# Remote magnetomechanical neuromodulation uncovers a novel therapeutic mechanism for alleviating Parkinsonian symptoms in freely moving mice

**DOI:** 10.1101/2025.08.21.671452

**Authors:** Anouk Wolters, Lorenzo Signorelli, Christian Herff, Sophia Gimple, Renzo Riemens, Gunter Kenis, Kim Rijkers, Jyh-Jang Sun, Yasin Temel, Hans Clusmann, Danijela Gregurec, Sarah-Anna Hescham

**Affiliations:** Department of Neurosurgery, Mental Health and Neuroscience Research Institute, Maastricht University, Maastricht, the Netherlands; European Graduate School of Neuroscience (EURON), Maastricht University, Maastricht, the Netherlands; Department of Chemistry and Pharmacy, FAU Erlangen-Nuremberg, Erlangen, Germany; Department of Psychiatry and Neuropsychology, Mental Health and Neuroscience Research Institute (MHeNs), Maastricht University Medical Centre, Maastricht, the Netherlands; Academic center for epileptology Maastricht/Heeze, The Netherlands; Centre for Integrative Neuroscience (CIN), Maastricht University, Maastricht, the Netherlands; ATLAS Neuroengineering, Leuven, Belgium; College of Semiconductor Technology, Chung Yuan Christian University, Taoyuan City, Taiwan; Research Group Experimental Oto-rhino-laryngology, Department of Neurosciences, KU Leuven, Leuven, Belgium; Department of Neurosurgery, RWTH Aachen University Hospital, Aachen, Germany

**Keywords:** nanotechnology, magnetomechanical neuromodulation, DBS, nanoparticles, Piezo1, TRPV4

## Abstract

To overcome the limitations of invasive neuromodulation systems, we introduce a wireless magnetomechanical approach for remote, minimally invasive deep brain stimulation (DBS) without chronically implanted electrodes. This method leverages biocompatible nanoscale magnetite nanodiscs (MNDs) with ground vortex magnetisation, which undergo in-plane transitions under low-frequency alternating magnetic fields, thereby generating localised piconewton-scale torques. These torques engage endogenous mechanosensory pathways to modulate neural activity, enabling reversible stimulation without the need for genetic modifications. Calcium imaging validated the rapid neuromodulatory effects of MNDs *in vitro* and *ex vivo*, which motivated the subsequent application of magnetomechanical DBS to the subthalamic nucleus in mice. We demonstrated the remote control of motor behaviour in wild-type mice and significant restoration of motor function in a severe hemiparkinsonian model. This study established the first wireless therapeutic magnetomechanical neuromodulation platform that leverages biocompatible nanomaterials and endogenous mechanosensory ion channels, representing a promising step toward untethered, clinically translatable neurotechnology.

## Main

Deep brain stimulation (DBS) is a clinically established, invasive neuromodulation technique involving the surgical implantation of electrodes to deliver electrical stimulation to targeted deep brain regions. It has demonstrated significant therapeutic efficacy in neurological disorders, such as Parkinson’s disease (PD), essential tremor, and dystonia. Despite clinical success, the core DBS technology has remained largely unchanged since its initial implementation (1, 2). Reliance on permanently implanted hardware poses challenges related to surgical complexity, device longevity, and patient acceptance (3). Moreover, conventional DBS often lacks the spatial precision required to selectively target specific neural circuits because of the large volume of tissue activated by the standard electrodes. This can result in off-target effects, including unwanted motor or cognitive side effects, and contribute to variability in clinical outcomes (4). These limitations have motivated efforts to develop alternative neuromodulation strategies that preserve the effectiveness of DBS while reducing its invasiveness (5, 6).

Magnetic nanomaterials, which transduce magnetic fields into biological effects, are emerging as promising tools in bioelectronics and neuromodulation (7). Their ability to enable remote, localised stimulation without physical wiring makes them attractive for minimally invasive approaches. Magnetoelectric nanoparticles, for example, convert magnetic field input into electrical signals (8, 9). However, they often contain non-biocompatible elements with the risk of long-term degradation and present challenges in achieving consistent synthesis and control over their physical properties (10, 11). Magnetothermal approaches use magnetic nanoparticles to dissipate heat under an alternating magnetic field (AMF), thereby activating thermosensitive ion channels (12, 13). While effective, this approach offers limited temporal precision, relies on high concentrations of nanoparticles, raises concerns about the long-term effects of repeated neuronal heating, and typically relies on genetic modification to sensitise target neurons to thermal stimuli (14, 15).

Here, we introduce magnetomechanical DBS (mDBS; Fig. 1A), a wireless and minimally invasive neuromodulation method that employs our previously pioneered biocompatible magnetite nanodiscs (MNDs; Fig 1C and D) (16) to activate mechanosensitive ion channels in the brain, such as Piezo-type mechanosensitive ion channel component 1 (Piezo1) and transient receptor potential vanilloid 4 (TRPV4) (17, 18). When exposed to low-frequency AMF, MNDs generate piconewton torque (16), enabling the spatially localised mechanical stimulation of mechanosensitive neural cells (Fig. 1A).

**Figure 1.**
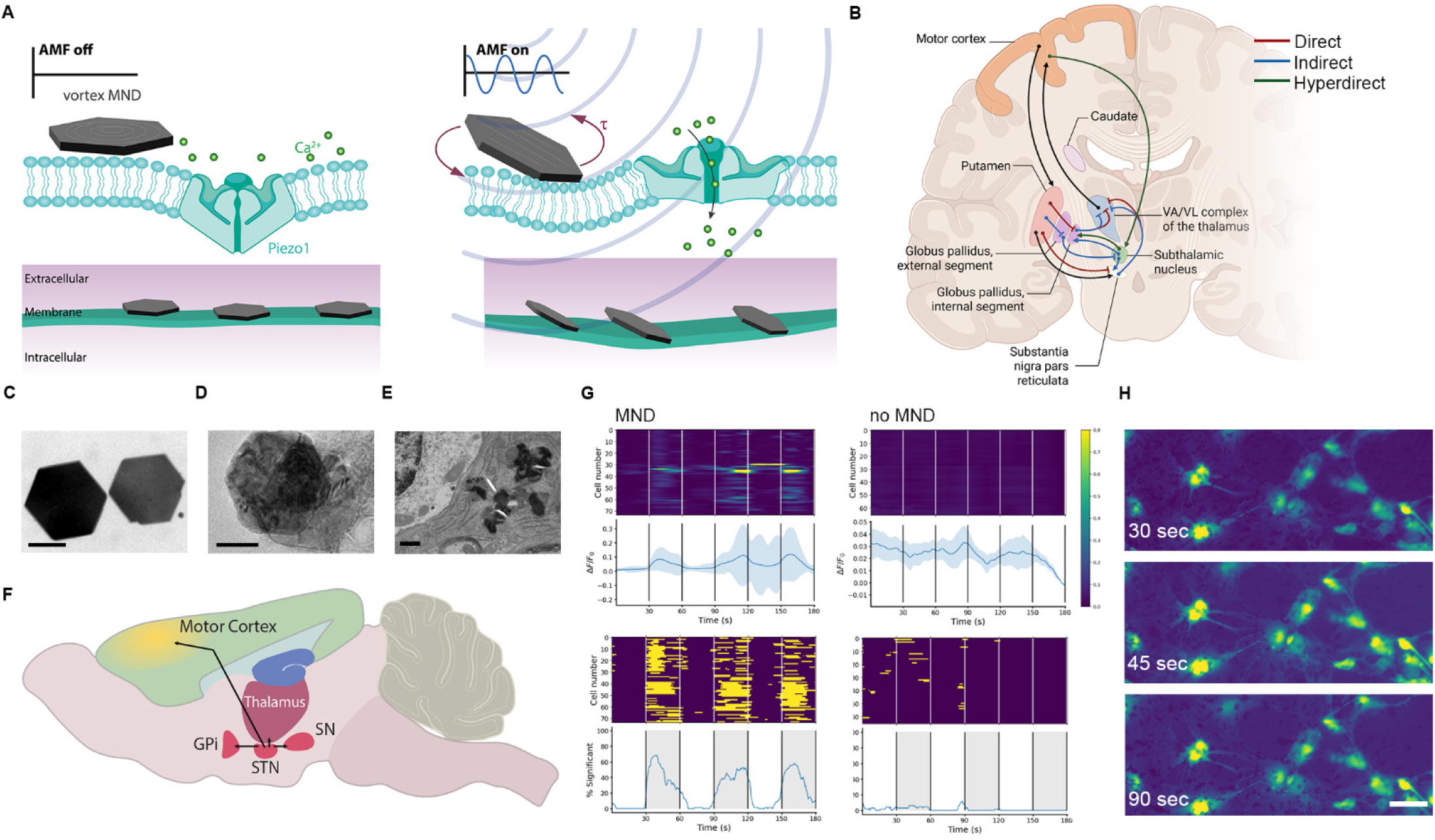
MNDs for magnetomechanical neuromodulation. **A.** An illustration of neuromodulation mediated by MNDs. **B.** Schematic illustration of the direct, indirect and hyperdirect pathways in the physiological condition. In the physiological condition, dopamine from the Substantia Nigra pars compacta (SNc) activates the direct pathway (red lines) and simultaneously inhibits the indirect pathway (blue lines). In addition, direct projections from the cortex are sent to the subthalamic nucleus (STN), which in turn sends them directly toward the Globus pallidus internus (GPi; green lines). The output structures, GPi and Substantia Nigra pars reticulata (SNr), which send efferent projections completing the cortico-basal ganglia-thalamo-cortical pathway (adapted from ref. (30)). **C.** Transmission electron microscopy (TEM) micrograph showing the morphology of MNDs synthesised via hydrothermal synthesis (scale bar = 100 nm). **D.** TEM micrograph showing poly(maleic anhydride-alt-1-octadecene-coated MNDs (scale bar = 100 nm). **E.** TEM micrograph showing MND within the STN of a mouse (scale bar = 100 nm). **F.** An illustration of a mouse brain highlighting the STN (pink) and showing the investigated regions being the primary motor cortex (yellow), SNr (pink), ventral lateral and ventral anterior complex of the thalamus (pink). **G.** Heatmaps (top) and significance plots (bottom) of fluorescence intensity changes recorded from Fluo-4 AM transients observed in Piezo1 and TRPV4 expressing human embryonic stem cell (hESC)-derived neurons during magnetic field stimulus with (left) and without (right) MNDs. Each condition was repeated across six different cultures, with ∼10 cells randomly selected for analysis. **H.** False colour stills from a representative video of hESC-derived neurons (scale bar = 50 µm).

A recent study reported magnetomechanical stimulation using MNDs (19), building directly on our reported MND design and magnetic parameter optimisation. However, the use of higher frequency and higher amplitude magnetic fields, along with a focus on transient receptor potential cation channels, may have contributed to limited efficacy, potentially affecting both specificity and translational relevance (16). In contrast, strategies such as mTorquer require large concentrations (50 mg/ml) coupled with viral transduction to overexpress Piezo1 for functional efficacy, which further complicates its clinical translation (20).

Our work targets well-characterised endogenous mechanosensitive channels such as Piezo1 and TRPV4, which are expressed in neurons and glial cells in multiple regions of the mammalian central nervous system (CNS) (21, 22). These channels play a key role in transducing mechanical strain into electrical or biochemical responses (23, 24). Although their role in glia and vascular mechanotransduction in the CNS is well established (21, 25, 26), their contribution to neural circuit dynamics remains largely unexplored. Here, we harness these endogenously expressed mechanosensitive ion channels to enable wireless and spatially precise neuromodulation using MNDs actuated by clinically relevant low-frequency AMF parameters. We demonstrated that µg/ml concentrations of MNDs were sufficient to evoke robust neuromodulatory responses *in vitro*, *ex vivo*, and at µg doses in freely moving mice. This approach reveals a functional role for endogenous mechanosensitivity in modulating neural activity and provides a genetically unmodified approach for minimally invasive neuromodulation of deep brain circuits.

Crucially, we demonstrated functional recovery in a severe Parkinsonian mouse model, representing the first therapeutic application of fully endogenous magnetomechanical neuromodulation. Our mDBS approach may offer potential for future human applications, with the possibility of improving patient adherence and clinical outcomes in DBS treatment (13).

## Results

### MNDs mediate calcium influx *in vitro* via endogenous mechanosensitive ion channels

We examined whether MNDs can induce intracellular calcium responses via magnetomechanical stimulation, and thus, the ability of MNDs to stimulate mechanosensitive ion channels *in vitro*. Human embryonic stem cell (hESC)-derived neurons (H9 lineage, differentiated for >10 weeks) were incubated with the intracellular calcium indicator Fluo-4 AM and exposed to AMF stimulation (5 Hz, 28 mT) (16). Cells were treated with or without 30 µg/ml MNDs and subjected to three 30s AMF On/AMF Off cycles using a custom-built solenoid coil (Extended Data Fig. 1A and C).

In cultures preincubated with MNDs, AMF pulses consistently showed robust increases in Fluo-4 fluorescence (ΔF/F_0_) (Fig. 1G and H). No response was observed in MND-free controls (Fig. 1G), MND-treated cultures that were not exposed to AMF (Extended Data Fig. S2C and D) or in the presence of the mechanosensitive cation channel inhibitor GsMTx4 (Extended Data Fig. S2E). Addition of the Piezo1 agonist Yoda1 also led to an increase in Fluo-4 fluorescence (Extended Data Fig. S2F), indicating that intracellular Ca^2+^ influx is mediated by mechanosensitive channels (27–29). The safety of magnetomechanical actuation under these conditions was confirmed by the absence of propidium iodide uptake 30 minutes after AMF stimulation (Extended Data Fig. S2G). Immunohistochemistry confirmed the expression of Piezo1 and TRPV4 in differentiated hESC-derived neurons (Extended Data Fig. S2H).

To assess the initial biocompatibility of the MNDs, we monitored HEK293 cell proliferation using phase-object confluence measurements. Over five days, untreated HEK293 cells showed a steady increase in confluence, reaching approximately an eight-fold increase (Extended Data Fig. S2A and B). Exposure to MND at concentrations of 15 µg/ml and 30 µg/ml did not inhibit cell growth, indicating the absence of cytotoxic effects. These findings suggest that MNDs do not impair HEK293 cell viability and proliferation at low concentrations. The concentration used for all further *in vitro* experiments was 30 µg/ml.

### MND-mediated magnetic stimulation evokes neuronal activation *ex vivo*

To address magnetomechanical activation in brain tissue, we incubated human organotypic brain slices (hBSCs) 3 days post-resection in artificial cerebrospinal fluid (aCSF) with ∼4 µg MNDs or without MNDs. Magnetomechanical stimulation was performed using the same custom-built solenoid coil system (Extended Data Fig. S1A and C) with three 20s AMF On/ Off cycles. Calcium imaging revealed that only neurons treated with MNDs exhibited a reproducible increase in calcium signals during the field-On epochs (Fig. 2A and Extended Data Fig. S3), confirming the effective mechanical stimulation in a physiologically relevant *ex vivo* environment. Immunohistochemistry confirmed the expression of Piezo1 and TRPV4 in human neurons (Fig. 2B).

**Figure 2.**
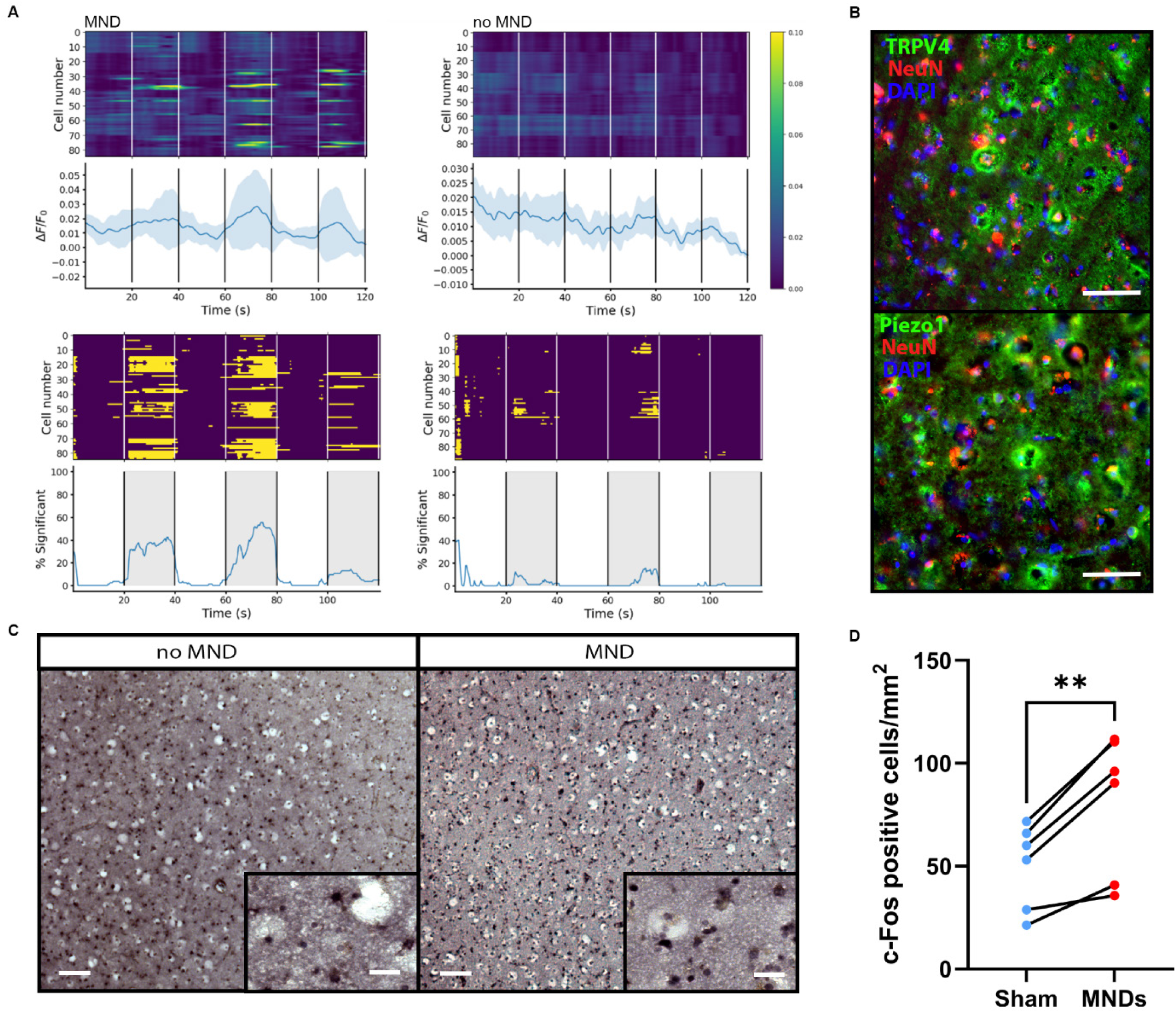
Ex vivo magnetomechanical control. **A.** Heatmaps (top) and significance plots (bottom) of fluorescence intensity changes recorded from Fluo-4 AM transients observed in Piezo1 and TRPV4 expressing hBSC during magnetic field stimulus with (left) and without (right) MNDs. Each condition was repeated across six different cultures, with ∼15 cells randomly selected for analysis. **B.** Representative high-power photomicrographs of hBSC stained for NeuN and TRPV4 or Piezo1 (scale bar = 50 µm). **C.** Representative low-power photomicrographs (scale bar = 100 µm) of hBSC stained for c-Fos (K-25), each with a high-power photomicrograph inset in the lower corner (scale bar = 25 µm) in slices receiving mDBS (n=6) and controls (n=6). **D.** Comparisons were made between AMF only (n=6) and mDBS (n=6) slices. mDBS-treated slices showed significantly increased c-Fos (K-25) expression (p = 0.0018). ** Indicates p < 0.01. Data presented as mean ± S.E.M., one-tailed paired t-test.

To determine whether MND-mediated stimulation activates functional neural circuits, tissues were fixed 90 minutes after AMF treatment and subsequently immunolabeled for c-Fos, a marker of recent neuronal activity (31). Slices incubated with MNDs and exposed to AMF showed a significant increase in c-Fos expression compared to AMF only (t(5)=5.4180, p = 0.0018, Fig. 2C and D). These findings demonstrate that magnetomechanical actuation can effectively induce activity-dependent gene expression in intact human brain tissues.

### *In vivo* MND-based magnetomechanical stimulation modulates motor behaviour in mice

Next, we tested the ability of MNDs to modulate brain activity *in vivo*. Adult C57BL/6J mice received unilateral stereotaxic injections of 1.5 µg MNDs (1.5 µl per animal) into the subthalamic nucleus (STN) (Fig. 1E), a deep brain structure clinically targeted for DBS in PD. Unilateral targeting was chosen based on evidence that classical unilateral STN DBS induces changes in motor behaviour in rodents (32). Motor behaviour in response to magnetomechanical STN stimulation was evaluated 1-2 weeks post-injection.

To compare AMF On and Off conditions, mice were placed inside the AMF coil for 3 minutes at 5 Hz and 28 mT. AMF stimulation led to significant improvement in motor performance on the rotarod task, which evaluates balance, coordination, physical condition, and motor planning. Mice exposed to AMF remained on the rotating rod for significantly longer than during the AMF Off sessions (t(9)=2.708, p = 0.0120, Fig. 4B). In addition, mice were evaluated using the open field test (OFT) for 5 minutes. The OFT assesses general locomotor behaviour in a novel open arena. Following AMF On stimulation, mice exhibited significantly reduced movement compared to the AMF Off condition (t(10)=1.860, p = 0.0463; Extended Data Fig. S4A-C). In contrast,

The rotational behaviour was evaluated in a circular plexiglass arena (Ø 9 mm, height 30 cm) fitted within the custom-built solenoid coil (Extended Data Fig. S1D). Mice showed no significant change in contralateral rotations (t(20)=1.2209, n.s., Extended Data Fig. S4D and E) or ipsilateral rotations (t(20)=0.2868, n.s., Extended Data Fig. S4D and E) during the AMF On versus AMF Off conditions. While a trend toward reduced ipsilateral rotations was observed during the AMF On condition, the overall rotational symmetry remained unchanged.

AMF stimulation also produced significant improvements in gait- and balance-related general coordination, as assessed using the CatWalk XT test (33). Mice exhibited decreased run duration (p = 0.0173, Fig. 3C), no change in the number of steps (Fig. 3D), increased support of the diagonal (p = 0.0275, Fig. 3E), and increased stride length (p ≤ 0.0346, Fig. 3F). Additional changes included a decreased print area (p = 0.0176, Fig. 3G), decreased initial dual stance (p ≤ 0.0276, Fig. 3H), and lower body speed variation (p ≤ 0.0290, Fig. 3I).

**Figure 3.**
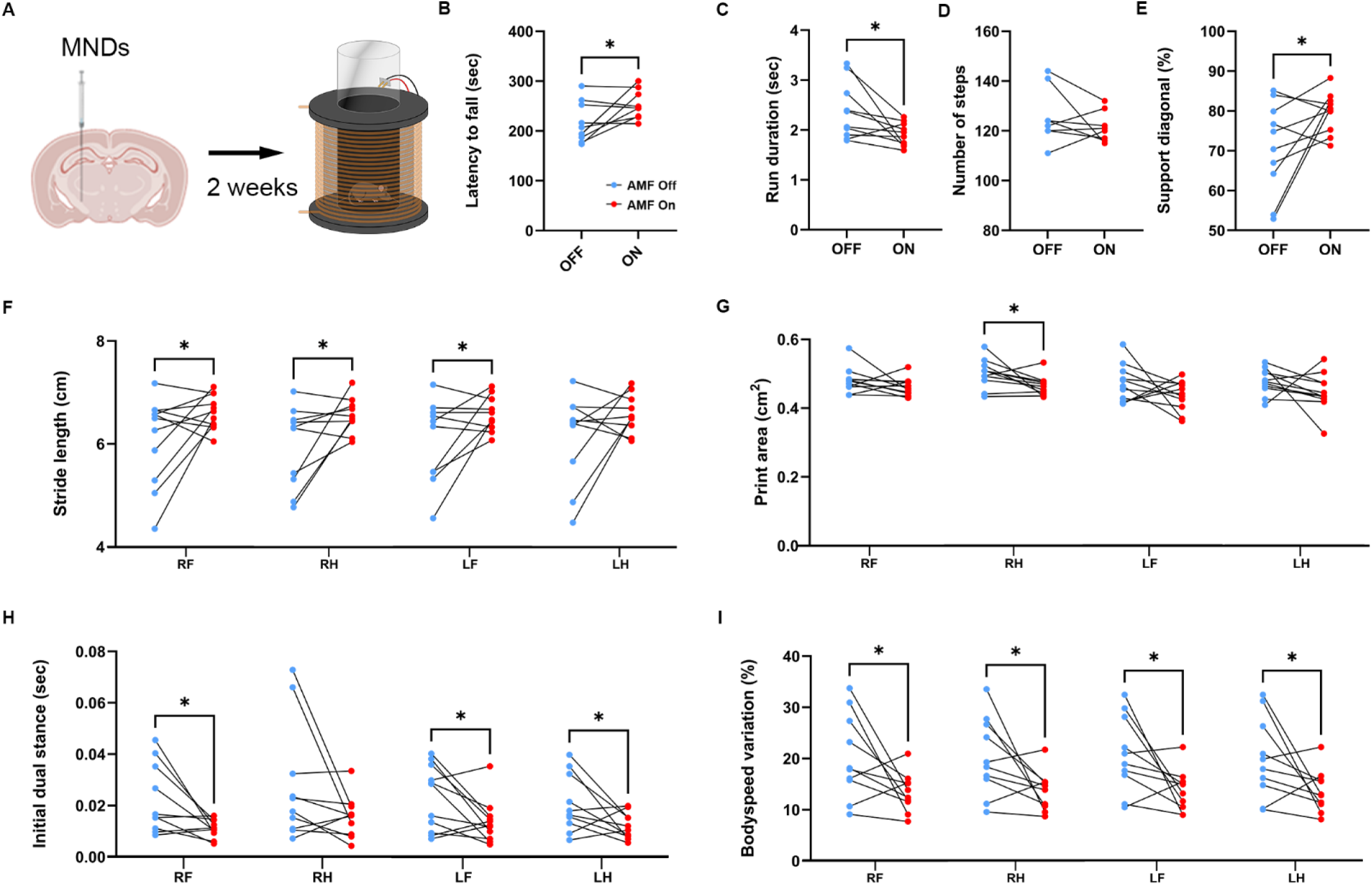
Magnetomechanical STN DBS improves motor behaviour in naïve male mice. **A**. In vivo experimental scheme. **B**. Latency to fall from the rotarod (in seconds) during AMF Off and AMF On conditions averaged from four consecutive trials. Mice stimulated in an AMF 3 min before the rotarod significantly improved their motor performance (paired t-test, t(9)=2.708, p = 0.0120). **C-I** Representative CatWalk XT results of five runs. AMF stimulation resulted in faster run duration (p = 0.0173) **(C)**, no change in number of steps (n.s) **(D)**, increased diagonal limb support (p = 0.0275) **(E)**, increased stride length (p ≤ 0.0346) **(F)**, decreased paw print area (p = 0.0176) **(G)**, decreased initial dual stance (p ≤ 0.0276) **(H)**, and reduced body speed variation (p ≤ 0.0290) **(I)** of STN mDBS (n=11) during AMF On and Off conditions as revealed by paired t-test. * Indicates p < 0.05. RF: right front paw, RH: right hind paw, LF: left front paw, LH: left hind paw.

Supplementary analyses revealed a decreased maximum contact area (p ≤ 0.0255) (Extended Data Fig. S7A), decreased terminal dual stance (p ≤ 0.0425) (Extended Data Fig. S7B), decreased print length (p = 0.0026) (Extended Data Fig. S7C), no change in print width (Extended Data S7D), no change in speed duty cycle (Extended Data S7E), run max variation (p = 0.0075) (Extended Data Fig. S7F), and decreased support three (p = 0.0134) (Extended Data Fig. S7G), indicating an enhanced dynamic coordination and postural control following AMF stimulation.

### Therapeutic efficacy of magnetomechanical DBS in a severe Parkinsonian mouse model

To assess the clinical relevance of mDBS, we applied this approach to a well-established model of PD, namely mice with a unilateral 6-hydroxydopamine (6-OHDA) lesion targeting the nigrostriatal pathway in the STN. Given that STN stimulation is a standard DBS target for levodopa-resistant motor complications, MNDs were unilaterally injected into the STN of hemiparkinsonian mice. This hemiparkinsonian model is characterised by selective dopaminergic neuron loss on one side of the brain, mimicking the motor deficits observed in PD (34). We found that a unilateral 6-OHDA injection resulted in a 90% reduction in Tyrosine Hydroxylase (TH) expression in the ipsilateral substantia nigra pars compacta (SNc) when compared to the contralateral hemisphere (p < 0.0001) (Fig. 4B). To compare the behavioural effects of mDBS, sham and MND-treated mice were evaluated for their performance on the rotarod. Performance was significantly improved in the MND group compared to the sham control group (t(19)=1.875, p = 0.0381) (Fig. 4C), indicating improved balance, coordination, physical condition, and motor planning under magnetomechanical stimulation.

**Figure 4.**
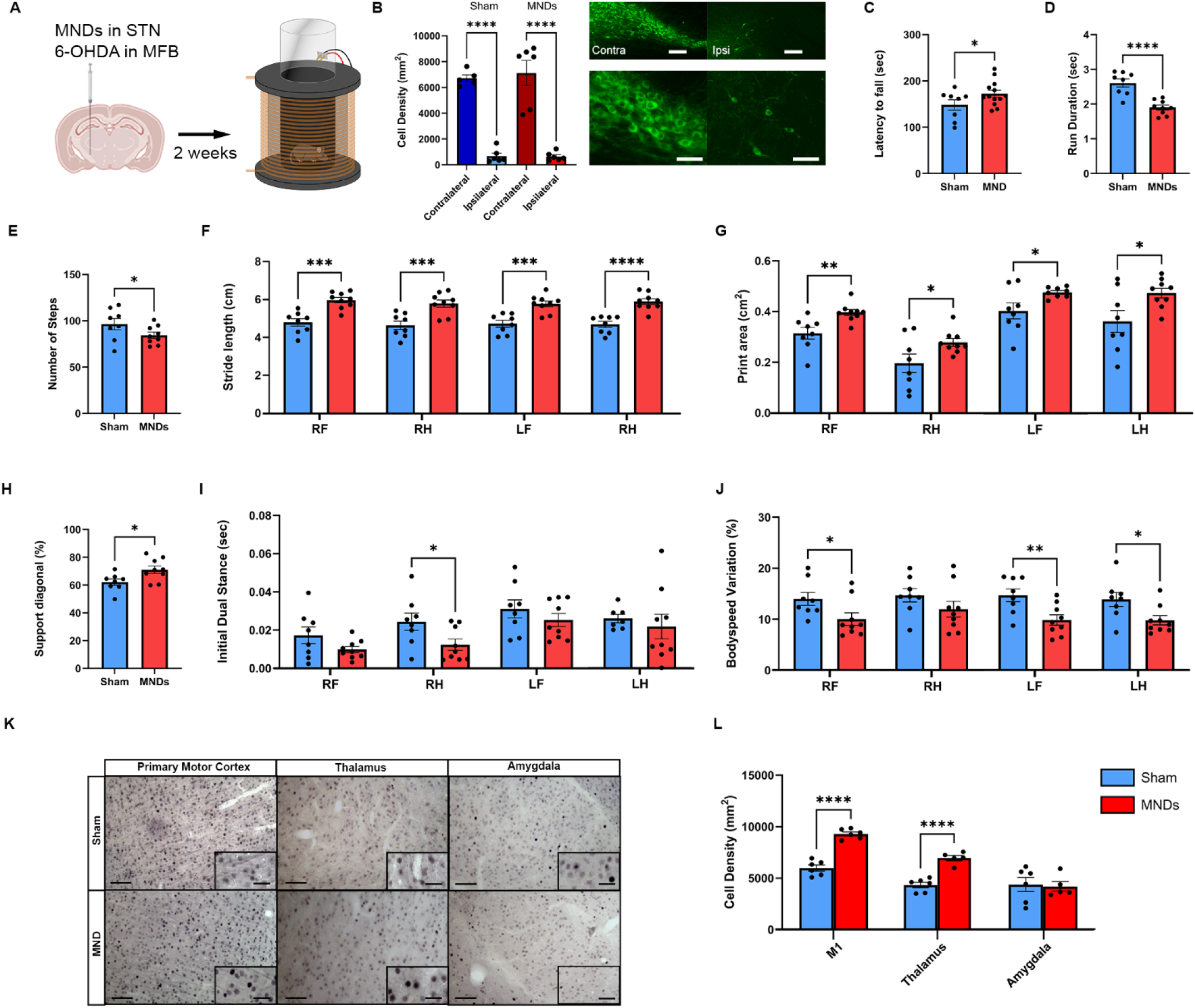
magnetomechanical STN DBS alleviates severe Parkinsonian symptoms. **A.** In vivo experimental scheme for the translational study. **B.** Representative low-power (scale bar = 1000 µm) and high-power (scale bar = 50 µm) photomicrographs of Tyrosine Hydroxylase (TH)-positive cells in the substantia nigra pars compacta (SNc) contralateral and ipsilateral to the 6-OHDA lesion. 6-OHDA lesion resulted in a 90% reduction in TH levels on the ipsilateral SNc when compared to the contralateral side. Data presented as mean ± S.E.M., one-way ANOVA, p < 0.0001. **C**. Latency to fall from the rotarod (in seconds) between the sham and MNDs group averaged from five consecutive trials on 2 test days. Mice stimulated in the MNDs group showed a significantly improved motor performance during the rotarod (t(19)=1.875, p = 0.0381). **D–J** Representative CatWalk XT results of four runs. AMF stimulation resulted in faster run duration (p < 0.0001) **(D)**, less number of steps (p = 0.0472) **(E)**, increased stride length (p ≤ 0.0006) **(F)**, decreased paw print area (p ≤ 0.0224) **(G)**, increased diagonal limb support (p = 0.0118) **(H)**, decreased initial dual stance (p = 0.0190) **(I)**, and decreased body speed variation (p = 0.0352) **(J)** between sham (n=9) and STN mDBS (n=8) as revealed by an independent one-tailed t-test. **K.** Representative low-power photomicrographs (scale bar = 500 µm) of coronal brain sections stained for c-Fos (K-25) showing the primary motor cortex (M1), ventral anterior and ventral lateral complex of the thalamus, and amygdala each with high-power photomicrograph inserts in the lower right corner (scale bar = 50 µm) in mice receiving sham (n=6) and STN mDBS (n=6) **L.** Comparisons were made between sham mice (n=6) and STN mDBS (n=6). STN mDBS animals showed increased c-Fos expression in motor areas such as M1 (P < 0.0001) and the ventral anterior and ventral lateral complex of the thalamus (P < 0.0001), whereas the amygdala, a non-motor area, was not affected. * Indicates p < 0.05, ** P < 0.01, *** p < 0.001, **** p < 0.0001. Data presented as mean ± S.E.M. RF: right front paw, RH: right hind paw, LF: left front paw, LH: left hind paw.

Rotational behaviour was quantified as previously described, with ipsilateral or contralateral rotations around the body axis. While a trend toward reduced ipsilateral turning was observed in MND-treated mice, statistical analyses revealed no significant differences in ipsilateral (t(20)=0.2868, n.s.) (Extended Data Fig. S5A) or contralateral rotations (t(20)=0.2868, n.s.) (Extended Data Fig. S5B) between sham and STN mDBS. However, the overall distribution of the rotational behaviour appeared to be more balanced under stimulation, suggesting subtle modulatory effects.

Additionally, sham and MND-treated mice were evaluated in the OFT for 5 minutes as previously described. No significant difference in the total distance travelled was observed between the MND and sham groups (t(15)=0.1679, n.s.) (Extended Data Fig. S6A), although a positive trend toward increased locomotion was observed in the MND group. Similarly, the average velocity did not differ between groups (t(14)=0.9838, n.s.) (Extended Data Fig. S6B). The time spent in the centre (t(15)=1.142, p = 0.1356) (Extended Data Fig. S6C), corners (t(15)=0.9593, n.s.) (Extended Data Fig. S6D), and along the walls (t(15)=0.08598, n.s.) (Extended Data Fig. S6E) of the OFT arena were also comparable between the groups, indicating no anxiety-related behaviour (Extended Data Fig. S6).

In the CatWalk XT test, AMF stimulation induced significant changes in multiple gait and balance-related parameters, reflecting improved coordination and motor control. MND-treated mice exhibited a decreased run duration (p < 0.0001) (Fig. 4D), reduced number of steps (p = 0.0472) (Fig. 54E), increased stride length (p ≤ 0.0006) (Fig. 4F), increased print area (p ≤ 0.0224) (Fig. 4G), and support of the diagonal (p = 0.0118) (Fig. 4H). The contact dynamics also changed with an elevated maximum contact area (p ≤ 0.0228) (Extended Data Fig. S8A), increased terminal dual stance (p ≤ 0.0246) (Extended Data Fig. S8B) and a higher initial dual stance (p = 0.0190) (Fig. 4I). Additional improvements included increased print dimensions, length (p ≤ 0.0110) (Extended Data Fig S8C), and width (p ≤ 0.0358) (Extended Data Fig. S8D). Measures of gait variability were reduced, including decreased body speed variation (p = 0.0352) (Fig. 4J), speed duty cycle (p = 0.0352) (Extended Data Fig. S8E), and maximum run variation (p = 0.0485) (Extended Data Fig. S8F).

Consistent with behavioural effects, magnetomechanical stimulation activated neural circuits across motor regions. In AMF-stimulated mice (n=6), a significantly higher proportion of c-Fos positive cells was detected in the motor cortex (t(10)=9.405, P < 0.0001) and ventrolateral and ventral anterior thalamic complex (t(9)=6.958, p < 0.0001) compared to the sham control group (n=6). No difference was observed in the amygdala (t(9)=0.2276, n.s.) in AMF-stimulated mice (n=5) when compared to the sham control group (n=6) (Fig. 4K and L), confirming the effective activation and recruitment of areas of the cortico-basal ganglia-thalamo-cortical circuitry (Fig. 1B and F).

### Magnetomechanical STN Stimulation Drives Network-Level Neural Responses *In Vivo*

To investigate the effects of low-frequency magnetomechanical activation of the STN on basal ganglia output and unravel network-level neural responses *in vivo*, we performed high-density extracellular recordings targeting the SNr and GPi in a hemi-parkinsonian mouse model (Fig. 5A). Spike sorting and assignment of single units were based on stereotactic coordinates and confirmed histologically. During electrophysiological recordings, repeated measures ANOVA revealed a significant effect of AMF stimulation on the firing rate of SNr neurons across baseline, during, and after stimulation (F(2,12) = 6.802, p = 0.0063) (Fig. 5B and C). Further post-hoc comparisons revealed a significant increase in firing rate between baseline and stimulation (p = 0.0430) and between baseline and post-stimulation (p = 0.0280). At the same time, no difference was found between stimulation and post-stimulation. Similar results were observed in the GPi, with repeated measures ANOVA indicating a significant effect across time points (F(2,5) = 14.25, p = 0.0071) (Fig. 5E and F). In particular, a significant increase in firing rate was observed between baseline and stimulation (p = 0.0302) and between baseline and post-stimulation (p = 0.0320), while no difference was found between stimulation and post-stimulation. These findings suggest that low-frequency magnetomechanical stimulation can modulate basal ganglia output in a manner comparable to conventional high-frequency STN-DBS (35), potentially preserving or restoring more physiological neural activity patterns within the network.

**Figure 5.**
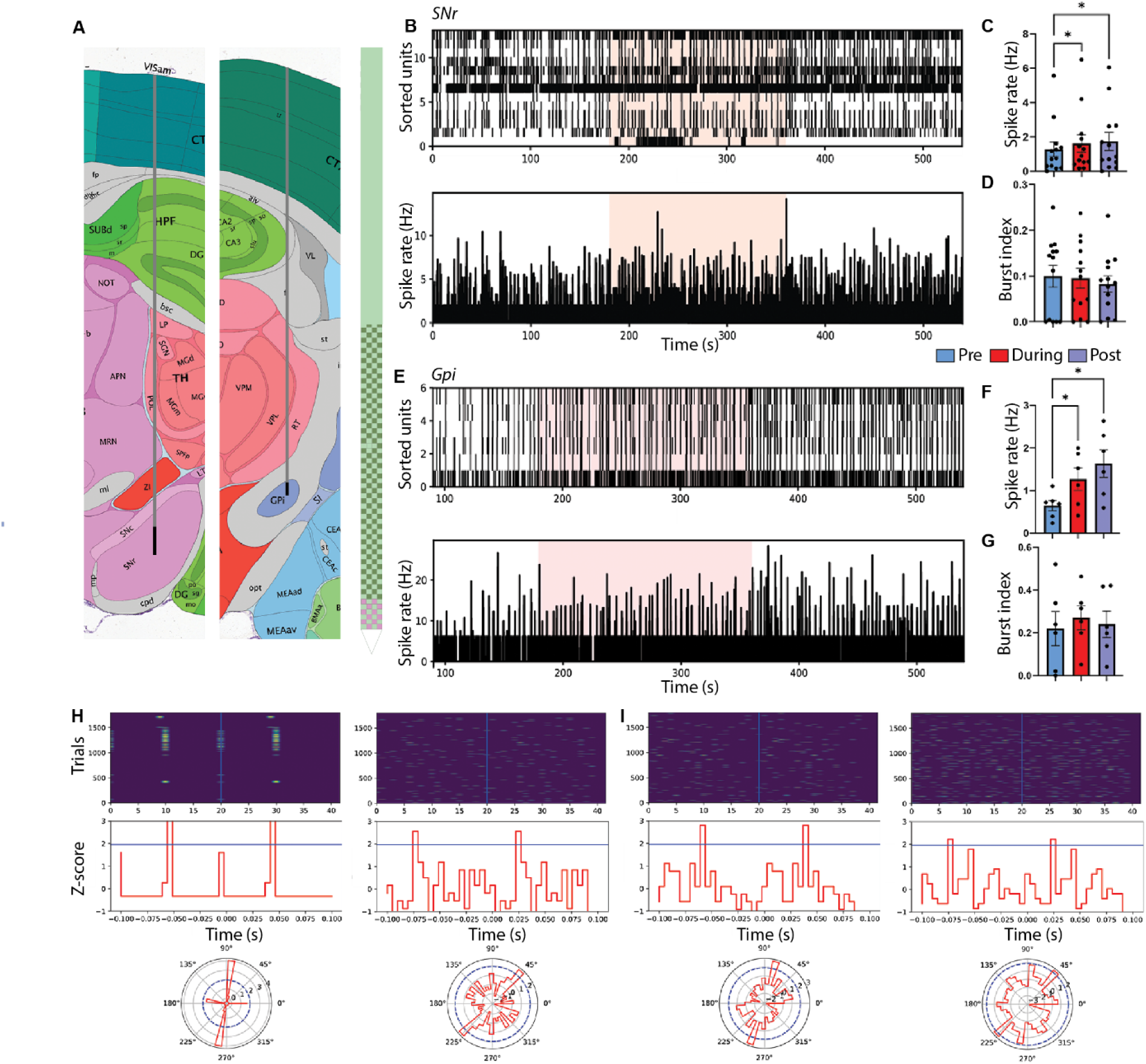
Magnetomechanical STN DBS neural responses. **A.** Neuropixels probe implantation trajectories to SNr and GPi according to the Allen Mouse atlas. The probes’ contact points within the target are displayed in pink (SNr) and blue (GPi). **B.** Raster plot (top) of 13 units recorded before, during and after AMF stimulation in the substantia nigra. Burst index (bottom) displayed as the summed spike rate of all units, binned at 50 ms intervals. **C.** Average spike rates of the 13 SNr units significantly increased (p < 0.05) during and after AMF stimulation compared to the pre-stimulation baseline. **D.** The average burst index remained unchanged during and after AMF stimulation relative to the pre-stimulation period. **E.** Raster plot (top) of 6 units recorded before, during and after AMF stimulation in the globus pallidus internus. Burst index (bottom) displayed as the summed spike rate of all units, binned at 50 ms intervals. **F.** Average spike rates of the 6 GPi units significantly increased (p < 0.05) during and after AMF stimulation compared to the pre-stimulation baseline. **G.** The average burst index remained stable during and after AMF stimulation relative to the pre-stimulation period. **H.** Two representative substantia nigra units responding to AMF stimulation. Top: Raster plots of all trials aligned to single AMF stimulation events, showing spike activity 0.1 seconds before and after stimulation. Bottom: Polar plots indicating preferred phase of AMF stimulation. The blue line denotes a Z-score threshold of 1.96. **I.** Two representative globus pallidus internus units responding to AMF stimulation. * Indicates p < 0.05. Data presented as mean ± S.E.M. Repeated-measures ANOVA with Bonferroni post hoc correction.

Interestingly, SNr and GPi neurons showed no changes in bursting activity during AMF stimulation (Fig. 5D and G). Bursting activity in the basal ganglia nuclei is a well-established electrophysiological hallmark of PD, due to dopamine depletion and is typically associated with impaired motor function (36–38). Conventional high-frequency STN DBS suppresses bursting activity (38). Therefore, our findings suggest that AMF stimulation modulates neuronal firing rates without affecting pathological bursting patterns, indicating a distinct mechanism of action compared to conventional high-frequency DBS.

We next investigated whether neuronal activity in basal ganglia output structures was entrained by AMF stimulation. Representative single-unit recordings from the SNr revealed time-locked responses, with spikes aligned to stimulation onset showing changes in firing within a 100-ms window surrounding the event (Fig. 5H, top). Phase analysis further demonstrated significant locking of spikes to the AMF, with preferred phases exceeding the statistical threshold (Z= 1.96; Fig. 5H, bottom). Similarly, recordings from the GPi showed representative units that exhibited phase-locked responses to AMF stimulation (Fig. 5I).

## Discussion

There is an increasing interest in advanced neuromodulation technologies for PD, particularly in cases where pharmacological treatments are insufficient. STN-DBS has proven to be more effective for advanced PD, demonstrating increased efficacy over dopaminergic medication alone in managing motor symptoms and improving the quality of life in selected patients (39). The STN plays a central role in the cortico-basal ganglia-thalamo-cortical circuitry, exerting excitatory glutamatergic projections to downstream nuclei, including the globus pallidus and substantia nigra (40, 41). Given this functional importance, there is an increasing demand for neuromodulatory approaches that offer greater specificity, improved safety and long-term stability, while minimising invasiveness. Recent innovations in miniaturised and wireless systems have sought to address these limitations by enabling remote neural control without the need for chronically implanted hardware or frequent surgical interventions such as battery replacement (9, 13). Magnetothermal DBS has emerged as a promising strategy in this context, allowing remote neuromodulation through externally applied magnetic fields (13, 42, 43). However, to date, no clinical studies targeting neurological disorders have been reported. This is largely due to limitations such as reliance on transgenes, low temporal resolution and high nanoparticle concentrations (12, 13). Recently, approaches capable of delivering mechanical or electrical stimuli without requiring genetic modification have drawn attention because of their translational potential (8, 9, 15, 16).

In this study, we introduce mDBS, a wireless approach using injectable, anisotropic MNDs with vortex magnetisation that transduce externally applied AMF energy into mechanical forces to activate neurons in the STN. After characterising the magnetisation behaviour and confirming previously demonstrated biocompatibility of MNDs stabilised with poly(maleic anhydride-alt-1-octadecene) (PMAO) and their non-toxic effects, we validated the mechanistic underpinnings in *in vitro* models.

Through Ca^2+^ imaging in hBSC and hESC-derived neurons, we observed that MND stimulation induced transient calcium influx through mechanosensitive ion channels, predominantly mediated by Piezo1. Pharmacological inhibition of Piezo1 and TRPV4 using GsMTx4 significantly reduced AMF-induced activity, whereas the Piezo1 agonist Yoda1 potentiated responses. These results confirm that MND-driven AMF exposure causes torques sufficient for membrane deformation and Piezo1 opening, initiating a calcium influx cascade, similar to the results of our reported MND application to activate dorsal root ganglions dominated by Piezo2, with ∼18 pN exerted from a single MND under 5 Hz and 28 mT AMFs (16).

Immunohistochemical analysis further revealed the presence of Piezo1 and TRPV4 in *ex vivo* brain tissue, suggesting the possibility of multi-channel mechanotransduction. Although our functional experiments focused on Piezo1, TRPV4 is known to respond to mechanical and osmotic stimuli and is involved in neurosensory and glial modulation (23, 44). Its expression raises the possibility of parallel or synergistic activation of pathways during mDBS. However, TRPV4 activation often requires more sustained or indirect stimuli compared to Piezo1’s direct bilayer tension gating (24). Future studies using TRPV4 antagonists or knockout models may reveal whether this channel contributes to neuromodulatory outcomes, particularly under chronic stimulation. Importantly, the presence of both channels highlights the potential of the mDBS platform to engage multiple endogenous pathways, thus enhancing its utility across diverse neurobiological contexts.

In naïve mice, unilateral stereotactic injection of MNDs into the STN, followed by AMF stimulation, significantly enhanced the performance in the rotarod and CatWalk XT assays. These findings suggest that mDBS can modulate motor circuits, even under baseline physiological conditions, thereby increasing motor coordination and balance. Notably, significant improvements in gait and balance assays, including walking speed, stride length, and variability in body speed, support the notion that magnetomechanical activation selectively enhances locomotor function, likely via motor-related neuronal pathways, including the STN and its downstream effects on the globus pallidus externa (GPe) and SNr via the indirect and hyperdirect pathways of the basal ganglia (45), as well as the motor thalamus (46), motor cortex (45), and brain stem centres, such as the pedunculopontine nucleus (47).

To test its therapeutic potential, mDBS was applied in a severe 6-OHDA hemiparkinsonian mouse model, which mimics advanced PD pathology with over 90% loss of TH-positive neurons in the SNc. While general locomotor activity in the OFT did not improve, likely because of the limited sensitivity of the test in detecting subtle motor improvements in severe dopaminergic depletion, mice treated with mDBS displayed significant benefits in rotarod latency and CatWalk XT parameters including increased walking speed, extended stride length, and a reduced duty cycle, suggesting a more efficient gait with less time spent in the stance phase. These results mirrored those observed in naïve animals.

Static CatWalk XT parameters also showed improvements relative to the sham controls. These behavioural outcomes are consistent with those of conventional STN DBS studies, including enhanced forelimb use asymmetry (48), enhanced treadmill performance (49), stepping and rotarod scores (50), and increased locomotor activity (51). Notably, static and dynamic Catwalk XT parameters have differential sensitivity to stimulation-induced changes in locomotor speed, suggesting that the dynamic improvement observed here reflects the selective engagement of fast locomotor circuits by mDBS.

To assess downstream neuronal activation, c-Fos immunostaining was used as a marker of neural activity (31). In hBSC, AMF stimulation in the presence of MNDs significantly increased c-Fos-positive cell counts compared to AMF alone. *In vivo*, a similar increase in c-Fos-positive cells was observed in the motor cortex and motor thalamus, confirming that MND-mediated mDBS can wirelessly drive cellular responses in mechanosensory deep brain structures. Although hBSCs offer a translational *ex vivo* platform, some limitations need to be considered, particularly the fact that this tissue is typically resected from pharmacoresistant epilepsy patients. Although it cannot entirely exclude the impact of disease-related changes, several factors support the broader relevance of our results. First, slices were obtained from cortical regions distant from the epileptogenic area, reducing epileptogenic effects. Second, our findings were confirmed *in vivo*, which are free from epileptic pathology, strengthening the generalisability of the observed neuromodulatory effects.

To examine the magnetomechanical activation of the STN, we performed high-density extracellular recordings targeting the SNr and GPi in a hemi-parkinsonian mouse model. AMF stimulation resulted in modulation of the basal ganglia output by increasing the firing rates in both structures. These increases were observed during and after stimulation, suggesting that mDBS modulates downstream basal ganglia output pathways. This effect is similar to the response to conventional high-frequency STN-DBS, which increases tonic firing in the SNr and GPi while disrupting pathological synchrony (35). Moreover, the increased excitability is in line with previous research indicating that 6-OHDA lesions disrupt the balance of synaptic inputs in favour of excitation, particularly in the GPi (52).

Despite the increase in firing rate, bursting activity in the SNr and GPi neurons remained unchanged during and after stimulation. Bursting is a hallmark of Parkinsonian basal ganglia activity and is closely related to dopamine depletion and motor dysfunction (36–38). While high-frequency DBS is known to suppress pathological bursting (38), MND-mediated stimulation did not alter this activity under the current parameters. This suggests that although magnetomechanical neuromodulation can modulate basal ganglia output, its mechanism of action may differ from that of conventional DBS. In addition, the ability of mDBS to impose phase-locked activity on basal ganglia output structures suggests temporally entrainment, which could be applied to disrupt pathological oscillations in Parkinsonian circuits (53). Together, these findings highlight the potential of mDBS to influence basal ganglia circuitry that is similar, but mechanistically distinct, from conventional DBS approaches (35, 40, 54).

In summary, this study presents a wireless, minimally invasive platform consisting of micromolar concentrations of biocompatible MNDs (1.5 µg per mouse) and simple, scalable solenoid coils for deep brain neuromodulation. By leveraging the magnetomechanical actuation of endogenous mechanosensitive ion channels dominated by Piezo1, mDBS enables targeted activation of the STN with high spatiotemporal resolution and without genetic tools or implanted hardware. Together, the behavioural and histological data support the conclusion that MND-mediated mDBS can effectively modulate neuronal activity in deep brain regions and induce functional changes. We propose that these effects are driven by selective activation of the thalamocortical circuitry, offering a minimally invasive, wireless neuromodulation strategy with therapeutic relevance for neurodegenerative diseases, such as PD.

## Materials & Methods

### Magnetic nanodiscs

The MNDs were synthesised through a two-step process. First, hematite nanodiscs were produced via hydrothermal synthesis. These were then reduced under hydrogen reflux at 360 °C while dispersed in triethanolamine, which served as a co-reducing agent, and oleic acid, employed as a dispersing agent. Oleic acid adsorbs onto the MNDs, conferring hydrophobicity and providing new binding sites for subsequent functionalisation with PMAO, applied at a concentration of 10 mg of PMAO per mg of MNDs, to enhance hydrophilicity, water dispersibility, and biocompatibility. When exposed to an AMF at a therapeutically relevant frequency of 5 Hz and an amplitude of 9 V_pp_, resulting in the generation of a magnetic field amplitude of 28 mT, sufficient to trigger the reversible activation of mechanosensitive ion channels through the torque produced by the MNDs.

### Cell and tissue culture

#### HEK293

HEK293 cells (ATCC® CRL-1573™) were cultured in EMEM (Gibco; 11095080) supplemented with 10% foetal bovine serum, maintained at 37 °C in a humidified atmosphere with 5% CO_2_. The cells were passaged every 3–4 days. Cells were seeded at 1.5 × 10^4^ cells/well in 96-well plates for toxicity analysis, and 1% penicillin-streptomycin was added to the cell culture medium.

#### H9 differentiated human neurons

H9 human embryonic stem cells (hESCs) obtained from WiCell were successfully expanded and confirmed to have a normal karyotype. The expression of pluripotency markers was validated following the protocol described by Marchetto *et al.* (55). H9 cells were cultured on Geltrex-coated plastic plates (Thermo Fisher Scientific) and maintained in E8 Flex medium (Thermo Fisher Scientific). Neural progenitor cells were generated using the STEMdiff™ Neural Induction Medium (Stemcell Technologies), according to the manufacturer’s instructions. Neural progenitor cells were maintained in STEMdiff Neural Progenitor Medium and subsequently matured into neurons using Brainphys medium, supplemented with growth factors.

In brief, hESCs were enzymatically dissociated using Gentle Dissociation Reagent (Stem Cell Technologies) and plated as single cells (3 x 10^6^ cells/ml) in AggreWell™ 800 microwell culture plates (Stem Cell Technologies) containing STEMdiff Neural Induction Medium, supplemented with SMAD inhibitors and y-27632, to generate EBs. Developing EBs were maintained in STEMdiff Neural Induction Medium supplemented with SMAD inhibitors for five days. After this period, the EBs were transferred to plates coated with polyornithine and laminin. Rosette-forming EBs were selectively isolated using an enzyme-free neural rosette selection reagent (Stem Cell Technologies) and plated on polyornithine-laminin-coated 35 mm dishes to generate neural progenitor cells (NPCs). These NPCs were cultured as high-density monolayers and seeded at low densities (4 × 10^4^ cells/cm²) for neuronal differentiation.

During neuronal differentiation, NPCs were cultured in neural differentiation medium supplemented with brain-derived neurotrophic factor (20 ng/ml, Peprotech), glial cell-derived neurotrophic factor (20 ng/ml, Peprotech), dibutyryl-cyclic AMP (1 mM, Sigma), ascorbic acid (200 nM, Sigma), and laminin (1 µg/ml) in BrainPhys Neuronal Medium (Stem Cell Technologies), which included N2 Supplement-A and SM1. The culture medium was refreshed twice a week throughout the ten-week differentiation process. Quality control was maintained to ensure that all cultures were free from mycoplasma contamination.

#### Human organotypic brain slice cultures

Slices for organotypic culture were prepared from the temporal cortex of eight patients (three male and five female, aged 25–63 years; Table 1) diagnosed with pharmacoresistant temporal lobe epilepsy. All patients provided written informed consent, and the protocols were approved by the Medical Ethics Review Committee (METC 2024-0496). Tissue processing followed a modified protocol based on the method described by Bak *et al*. (56). Tissue was transported from the operating room to the laboratory in slicing artificial cerebral spinal fluid (s-aCSF) composed of 110 mM choline chloride, 26 mM NaHCO₃, 1.25 mM Na₂HPO₄, 11.6 mM sodium ascorbate, 3.1 mM sodium pyruvate, 7 mM MgCl₂, 0.5 mM CaCl₂, 2.5 mM KCl, 10 mM glucose, and 1% penicillin/streptomycin/amphotericin B in demineralised water. The solution was carbonated with 95% oxygen and 5% CO₂ for at least 20 minutes before tissue resection. Under sterile conditions, the tissue was washed with carbonated s-aCSF, followed by careful dissection to remove meninges, capillaries, and damaged sections from the tissue block. Approximately 300 µm thick, Cortical tissue slices were cut using a vibratome (Leica VT1200, Leica Biosystems, Wetzlar, Germany), with each slice measuring approximately 5 × 5 mm. These slices were placed on 0.4 µm pore-size 6-well cell culture inserts (CellQART 9300414, SABEU, Germany), creating an air-liquid interface using intermediate step HEPES medium (ISHM).

**Table 1:** Patient demographics and experimental applications.

| **Gender** | **Age** | **Experiment(s)** |
| --- | --- | --- |
| *M* | 39 | IHC; C-Fos |
| *F* | 31 | IHC; C-Fos |
| *F* | 46 | IHC; C-Fos |
| *F* | 32 | IHC; C-Fos |
| *M* | 32 | IHC; C-Fos |
| *F* | 56 | IHC; C-Fos |
| *M* | 25 | Ca-Imaging |
| *F* | 63 | IHC; TRPV4 and Piezo1 |

ISHM was prepared by combining 23.5 ml DMEM F-12 and 23.5 ml Neurobasal medium, supplemented with 1 ml B27, 0.5 ml N2 supplement, 0.5 ml GlutaMAX, 0.5 ml penicillin/streptomycin, and 0.5 ml non-essential amino acids, yielding a total of 50 ml. To enhance the buffering capacity, 20 mM HEPES was added, and the mixture was stirred for at least 20 minutes. The slices were maintained at 37°C and 5% CO₂. After one hour, the slices on the inserts were transferred to a pre-equilibrated 6-well plate in an incubator containing artificial cerebrospinal fluid (aCSF). The aCSF composition included 125 mM NaCl, 25 mM NaHCO₃, 2.5 mM KCl, 1.25 mM NaH₂PO₄, 1 mM MgCl₂·6H₂O, 2 mM CaCl₂, 25 mM glucose, and 1% penicillin/streptomycin/amphotericin B in demineralised water. The aCSF was replaced two to three times per week. Tissue was afterwards collected in 4% paraformaldehyde (PFA) for 24 hours at 4°C. The tissue was then transferred to a 20% sucrose solution for a minimum of 24 hours. The tissue was then fresh-frozen and stored at -80°C.

#### Cell toxicity analysis

MNDs were suspended in cell culture medium at 15, 30, 60, and 120 µg/ml and then applied to the cells. Growth of HEK293 cells over 5 days post-MND administration using the IncuCyte® S3 imager (Sartorius, USA) with the basic analysis module. Phase area confluence values were normalised to the 0-hour time point for each condition and presented as a ratio to quantify relative changes over time.

### *In vitro* and *ex vivo* magnetomechanical stimulation

The magnetic stimulation setup was custom designed to fit into a Leica DIVI LFS microscope (Leica, Wetzlar, Germany) and accommodate a 35 mm dish. The system utilises a magnetic coil to generate an alternating current (AC) magnetic field focused along the centre of the cell culture well. A 1.32 mm-thick copper wire was wound around a plastic coil frame with 990 turns. This magnetic solenoid coil, used for the *in vivo* and *ex vivo* parts in this study, was custom-built in our lab by IDEE Research Engineering (for reference, see Gregurec et al. (16)). AC signals were produced by an Agilent 33210A 10 MHz Function/Arbitrary Waveform Generator and amplified using an Aetechron 7224 amplifier. The magnitude of the AC magnetic field was confirmed using a Wavecontrol SMP2 magnetometer. The coil used for in vitro experiments, custom-built by the Department of Chemistry and Pharmacy at the University of Friedrich-Alexander-Universität Erlangen-Nürnberg, was driven by a Crown DC-300 amplifier and an input signal from Keysight Technologies function generator.

In all the experiments, a 28 mT sine wave at 5 Hz was applied for 20–30 seconds during the 120–180 second recording period. Stimulating timings for differentiated H9 stem cells were as follows: 0–30 seconds with no field, 30–60 seconds with AMF, 60–90 seconds with no field, 90–120 seconds with AMF, 120–150 seconds with no field, and 150–180 seconds with AMF. The stimulation protocol for hBSCs was as follows: 0–20 seconds with no field, 20–40 seconds with an AC magnetic field (AMF), 40–60 seconds with no field, 60–80 seconds with AMF, 80–100 seconds with no field, and 100–120 seconds with AMF.

### Ca^2+^ transient experiments

#### Cell culture

Cells were loaded with 1 µM Fluo-4 AM dye (F14201, Invitrogen) and 1 µg/ml Propidium Iodide (P3566, Invitrogen) in phenol red-free Hank’s Balanced Salt Solution (HBSS;14025092, Gibco) for 30 minutes at 37°C. Following incubation, the cells were washed three times with HBSS, each lasting 5 minutes, by replacing half of the medium with fresh HBSS. After the final wash, half the HBSS was refreshed, and the cells designated for stimulation were incubated with MNDs for five additional minutes. Cells were then imaged using a Leica DIVI LFS microscope (Leica Biosystems, Wetzlar, Germany) with a 20x water-dipping objective for 3 minutes. Changes in fluorescence (ΔF/F_0_) were calculated using the procedure described in (Box 1, Jia *et al.* (*57*)). For the GsMTx4 condition, the cells were incubated with 1 µM GsMTx4 for 30 minutes post-wash. MNDs were added during the final 5 minutes of incubation. Imaging was conducted in the same manner as that for the other conditions. For the YODA1 condition, cells were treated similarly, but without the addition of MNDs. During imaging, 5 µM YODA1 was added after 30 seconds to assess Piezo1 reactivity. A maximum of six recordings were used, and approximately 10 cells per recording were analysed using the custom-written Python script based on the work of Jia *et al.* (57).

#### Human organotypic brain slice cultures

All procedures involving human tissue were conducted with the approval of the Ethics Committee of Maastricht University (METC 2024-0496). A day before the Ca^2+^ transient experiments, the tissue was separated to ensure that only one piece of tissue was in each transwell. On the morning of the experiments, the tissue was injected with 2 µl of a 2 mg/ml MND solution. Grids were placed on top of the tissue and left to settle for a minimum of 1.5 hours. Tissue was then loaded with a 1 µM Fluo-4 AM dye (F14201, Invitrogen) in aCSF for 30 minutes at 37 °C. After incubation with Fluo-4, the tissue was gently washed for 5 minutes with HBSS. After washing the tissue, the insert was removed and placed in a 35 mm dish. The insert was then filled with fresh HBSS and imaged, as previously described.

### Subjects

All procedures involving mice were conducted with the approval of the Animal Ethics Committee of Maastricht University (AVD10700202316737). The experiments were performed on 11 male naïve mice (C57BL/6J, Charles River), which were socially housed in a controlled environment maintained at 21 ± 1 °C with 40–60% humidity, following a reversed 12:12-hour light cycle (lights on at 21:00). All experimental manipulations were performed during the dark phase under red light, optimising conditions for rodent activity. At the time of surgery, the mice were aged 10–12 weeks and food and water were provided ad libitum. In the second cohort, 28 female wild-type mice (C57BL/6J, Charles River) were housed under identical conditions.

### Surgical procedure

Before anaesthesia, mice received a subcutaneous injection of buprenorphine at a dosage of 0.05 mg/kg as an analgesic. After 30 minutes, anaesthesia was induced and maintained using isoflurane at concentrations of 4% for induction and 1.5–3% for maintenance. Following adequate induction, each mouse was positioned in a stereotactic frame secured with ear bars and a mouse gas anaesthesia head holder. Body temperature was monitored and maintained at 37 °C using a thermoregulated heating pad. An ocular lubricant was applied to prevent drying of the eyes, and subcutaneous injection of 1% lidocaine was administered at the injection site for local anaesthesia and analgesia.

Subsequently, a burr hole was drilled above the left STN at coordinates AP: -2.06 mm, ML: -1.50 mm, DV: -4.50 mm. A microinjection apparatus was used to inject 1.5 µl of a 1 mg/ml MNDs solution at an infusion rate of 100 nl/min. Following the injection, the syringe needle was left in the brain for an additional 5 minutes before being slowly removed. In the second cohort, 1.5 µl of a 1 mg/ml MNDs solution was injected into the left STN, and 0.2 μl of 6-OHDA was unilaterally injected into the left median forebrain bundle (AP: −1.2 mm, ML: −1.1 mm, DV: −5 mm) at a rate of 100 nl/ min (3μg total). A total dropout rate of ∼10% was observed due to poor body condition of the 6-OHDA mice.

### *In vivo* magnetic stimulation

All *in vivo* magnetic stimulations were conducted using a custom coil system, which allowed the mice to move freely during the stimulation period. The system was designed to generate a 28 mT, 5 Hz alternating current (AC) magnetic field aligned with the central axis of the animal chamber. This magnetic field was produced by a single coil with the animal chamber positioned at its centre. A 1.32 mm-thick copper wire was wound around a plastic coil frame comprising 990 turns. An Agilent 33210A 10 MHz Function/Arbitrary Waveform Generator provided a 5 Hz sine wave that was amplified using an Aetechron 7224 power amplifier. The output was connected to an AC coil, and the magnetic field strength was verified using a magnetometer (Wavecontrol SMP2). For all stimulation experiments, the mice were exposed to a magnetic field for 3 minutes, with the coil either turned on or off.

### Behavioural tests

All behavioural tests were performed under dim light conditions, and animals were allowed to acclimate to the behavioural room for a minimum of 1 hour before the start of behavioural testing.

#### Rotational behaviour

The circular arena was made of plexiglass (9 cm diameter and 30 cm height, fitting within the magnetic coil). The mice were placed into the cylinder, and the AMF stimulation was turned on or off for 3 minutes. Two researchers manually scored the videos independently, and rotations were classified as ipsilateral or contralateral around the body axis. The frequency was assessed per 3 min and per 30-second time bins.

#### Open field test (OFT)

The open field test (OFT) consisted of a square arena with walls 25 cm high, measuring 40 cm in width and length. The behaviour tracking system was operated in semi-darkness for each mouse, employing a specialised tracking software (EthoVisionXT 15, Noldus Information Technology, Wageningen, The Netherlands). Each OFT lasted for 5 minutes, and each mouse was stimulated in an AMF coil for 3 minutes before testing.

#### Rotarod

A rotarod with the ability to accelerate using a grooved rotating beam (3 cm) raised 16 cm above the platform (model 47650, Ugo Basile, Italy) was used to measure the coordination of the mice. The latency to fall of the rotating rod was recorded. The mice were tested in four sessions, each lasting 300 seconds, with and without AMF stimulation, prior to the start. Testing was performed at speeds starting at 4 rpm and accelerated to 40 rpm within 300 seconds. Between trials, mice were allowed at least 2 min of rest to reduce stress and fatigue. Values are expressed as the mean latency to fall from the rotarod in all test trials.

#### Catwalk XT

The mice were assessed using the CatWalk-automated gait analysis system (CatWalk XT 10.6, Noldus). The apparatus consisted of a long glass walkway illuminated by fluorescent light directed toward the side of the glass floor. In a dimly lit setting, the light is reflected downwards, allowing a camera positioned beneath the glass to capture the footprints of the mouse as it moves along the walkway. The glass plate was cleaned and dried before testing each mouse to minimise the transmission of olfactory cues. In general, one successful test recording consisted of an average of four to five uninterrupted runs, with a comparable running speed and a maximum variation of 60%. The following parameters, including general, coordination, static, and dynamic aspects, were analysed to assess individual paw function and overall gait patterns: run duration, number of steps, number of step patterns, step pattern regularity index (%), stride length, max contact area, print area, print length, print width, initial dual stance, terminal dual stance and body speed variation, three limb support and diagonal limb support.

### Electrophysiological recordings

Recordings were performed in one anaesthetised, head-fixed, 6-OHDA mouse. The mouse was preoperatively injected with ketamine-medetomidine (K, 50 mg/kg; M, 0.2 mg/kg). Following adequate induction, the mouse was positioned in a custom-built stereotactic frame secured with ear bars and a head holder. Body temperature was monitored and maintained at 37 °C using a thermoregulated heating pad. Subsequently, a burr hole was drilled above the SNr at coordinates AP: -3.08 mm, ML: 1.5 mm, DV: 4.75 mm and GPi at coordinates AP: -1.34 mm, ML: 1.75 mm, DV: 4.55 mm. After the burr holes were drilled, the mouse was placed inside the AMF coil. The Neuropixels 1.0 probe was inserted inside the coil using a Sensapex uMP-3 NP micromanipulator and slowly inserted into the brain until the desired target was reached. Once the target was reached, the tissue was left to settle for 10-15 minutes per target. Recordings were acquired in three blocks: pre-stimulation, stimulation, and post-stimulation. Neural signals were acquired for the AP band at a sampling rate of 30 kHz, band-pass filtered between 0.3 kHz and 10 kHz, and an LFP band at a sampling rate of 2.5 kHz, band-pass filtered between 0.5 Hz and 500 Hz.

The recordings were spike-sorted using Kilosort4 (58) and manually curated using Phy (https://github.com/cortex-lab/phy) to identify single units. Single units were distinguished from multi-units by imposing a threshold of 5% in the contamination parameter computed by Phy, which is roughly the event rate ratio in the central 2 ms of the clusters’ autocorrelogram. The data were then imported into Python 3.8.0 (Anaconda 23.3.1) and analysed using custom-written scripts adapted from SpikeInterface (https://spikeinterface.readthedocs.io). The mechanical stability of the recordings, that is, the lack of significant drifting, was verified by visual inspection of drift maps, and dorsal-ventral positions of single units were aligned to the measured insertion depth of the probe.

Neurons with an average firing rate of <0.1 spikes/sec across the recording were excluded from the analysis. Spikes detected within 1 ms of stimulus cues were indissociable from the stimulation artefact and were therefore excluded from the study. To identify SNr neurons, we recorded a total of 13 single units from two separate recordings (n=1), and to study GPi neurons, we recorded a total of 6 single units from two separate recordings (n=1). Post-recording histological analysis was conducted to verify probe placement and assess any tissue damage, ensuring that recordings were made from the intended cortical regions.

### Tissue collection

At the end of the experiments, the mice were euthanised with an overdose of pentobarbital. Transcardial perfusion was performed using Tyrode’s buffer, followed by 4% PFA. Brains were then extracted and placed in fresh fixative for 24 hours at 4 °C. After fixation, the brains were transferred to 0.1% sodium azide (NaN₃) at 4 °C for long-term storage. For sectioning, the brains were embedded in 10% porcine skin gelatin (Sigma-Aldrich, Zwijndrecht, The Netherlands) and cut into 30 µm coronal sections using a vibratome (Leica®, Wetzlar, Germany). The sections were immediately placed in 0.1% NaN₃ and stored at 4 °C.

### Immunohistochemistry

#### Mouse brains

For immunohistochemistry, the sections were incubated overnight with polyclonal rabbit anti-c-Fos primary antibody (1:1000, Abcam, ab190289). The tissue was incubated with biotinylated donkey anti-rabbit secondary antibody (1:400, Invitrogen, Carlsbad, CA, USA) and avidin-biotin-peroxidase complex (1:800, Elite ABC kit, Vectastain, Burlingame, CA, USA). The staining was visualised using 3,3′-diaminobenzidine (DAB) combined with NiCl2 intensification. The slides were dehydrated and coverslipped using Entellan.

For immunofluorescence, the sections were incubated overnight with either polyclonal sheep anti-TH (1:2000, Abcam, ab113), polyclonal rabbit anti-IBA1 (1:2000, Invitrogen, PA5-27436), or biotin-conjugated rabbit anti-GFAP (1:2000, Merck, MAB3402B) primary antibody. Staining was visualised by immunofluorescence with donkey anti-sheep Alexa 488 (1:200), donkey anti-rabbit Alexa 594 (1:200), or streptavidin 594 (1:1000).

### Human organotypic brain slice cultures

#### C-Fos expression

For immunohistochemistry, the sections were incubated overnight with polyclonal rabbit anti-c-Fos primary antibody (1:1000, ab190289, Abcam). c-Fos immunohistochemistry included biotinylated donkey anti-rabbit secondary antibody (1:200, Invitrogen, Carlsbad, CA, USA) and avidin-biotin-peroxidase complex (1:400, Elite ABC kit, Vectastain, Burlingame, CA, USA). The staining was visualised using 3,3′-diaminobenzidine (DAB) combined with NiCl_2_ intensification.

#### TRPV4 and Piezo1 expression

For Immunofluorescence, hBSC sections were incubated overnight with polyclonal rabbit anti-TRPV4 (1:500, Alamone Labs, ACC-034) or for four nights with polyclonal rabbit anti-Piezo1 (1:100, Novus Biologicals, NBP1-78537) and monoclonal mouse anti-NeuN primary antibody (1:100, EMD Millipore, MAB377). TRPV4 and Piezo1 were visualised with donkey anti-rabbit Alexa 488 (1:200, Invitrogen, Carlsbad, CA, USA), and NeuN was visualised using donkey anti-mouse Alexa 594 (1:200, Invitrogen, Carlsbad, CA, USA). The nuclei were stained with Hoechst (1:10.000). Photomicrographs were taken using an Olympus Disk Spinning Unit (DSU) with a 20x objective.

### Stereology

The number of c-Fos- and TH-positive cells was counted using the stereological procedure optical fractionator. Counts were performed using a microscope (Olympus BX51W1), a motorised stage, and StereoInvestigator software (MicroBrightField, Williston, VT). All c- Fos- and TH-positive cells in an average of two to three sections per region of interest, 300 µm apart, were counted using a 20× objective. The total number of positive cells was estimated as a function of the number of cells counted and sampling probability.

The number of c-Fos-positive cells in hBSC tissue was counted using the same stereological procedure. All positively stained cells in an average of three sections were counted using a 20× objective. The total number of positive cells was estimated as a function of the number of cells counted and sampling probability.

### Immunocytochemistry

The cells were seeded at 2.0 × 10^5^ cells per well. Once the cells reached 80% confluence, the culture medium was removed, and the cells were washed with PBS. The cells were then fixed with 4% paraformaldehyde for 10 minutes. The cells were then washed again, incubated with 3% Triton-X for 10 minutes, and blocked for 30 minutes in 5% normal goat serum.

The fixed cells were incubated with polyclonal anti-Piezo1 and polyclonal anti-TRPV4 (1:200; Alomone Labs, Israel) primary antibodies for 50 minutes at room temperature. Next, the cells were incubated with donkey anti-rabbit Alexa 488 for Piezo1 or donkey anti-rabbit Alexa 594 for TRPV4 (1:200) secondary antibodies for 50 minutes at room temperature.

### Statistical analysis

All data are represented as mean ± standard error of the mean (S.E.M.). Statistical analyses were performed using GraphPad Prism software (version 10). Data normality and homogeneity of variance were checked using the Shapiro–Wilk test and normality plots. All behavioural data from the naïve experiment were analysed using a dependent one-tailed t-test to compare before and after AMF stimulation. Stereological cell counts were analysed using an independent samples t-test or one-way ANOVA. All behavioural data from 6-OHDA experiments were analysed using a one-tailed independent t-test. Electrophysiology data were analysed using a repeated-measures ANOVA with post hoc Bonferroni correction. Statistical significance was set at p < 0.05.

## Acknowledgements

We thank **Willine van de Wetering** and the Microscopy CORE Lab of M4I-FHML, Maastricht University, for their help with processing brains for TEM imaging, **Hossein Ghasem Damghani** for the help with 6-OHDA animal experiments and electrophysiology recordings, and **Geertjan van Zonneveld**, MUMC+, for the artwork. Finally, we wish to thank **Jyh-Jang Sun** from Atlas Neuro for his help with the electrophysiology analysis.

We also thank the ACE Epilepsy Study Group: Gwendolyn de Bruyn^1^, Albert Colon^1^, Jim Dings^1,2^, Marc Hendriks^1,3^, Danny Hilkman^1,4^, Christianne Hoeberigs^1,5^, Jochem van der Pol^5^, Lotte de Jong^1^, Kim Rijkers^1,2,6^, Sylvia Klinkenberg^1,7^, Vivianne van Kranen– Mastenbroek^1^, Jeske Nelissen^1,2,7^, Pieter Kubben^1,2^, Walter M. Palm^1,5^, Rob P.W. Rouhl^1,6,7^, Olaf Schijns^1,2,6^, Ilse van Straaten^1,8^, Simon Tousseyn^1,6^, Marielle Vlooswijk^1,7^, Louis Wagner^1^, and Dorien Weckhuysen^1^.

1 Academic Centre for Epileptology Kempenhaeghe/MUMC+

2 Department of Neurosurgery, MUMC+

3 Donders Institute for Brain, Cognition and Behaviour, Radboud University

4 Department of Clinical Neurophysiology, MUMC+

5 Department of Medical Imaging, MUMC+

6 School for Mental Health & Neuroscience Maastricht University

7 Department of Neurology, MUMC+

8 Department of Electrical Engineering, Eindhoven University of Technology

## Author Contributions

A.W., L.S., D.G, and S.H. designed the experiments and wrote the paper; D.G. and L.S. synthesised, functionalised and characterised the MNDs; A.W. and R.R. conducted the *in vitr*o work; D.G. designed the AMF coils for *in vivo* experiments; A.W. conducted data collection and analysis for all *in vitro, ex vivo* and *in vivo* experiments; S.H. performed the surgeries of all in vivo experiments; L.S. participated in naïve animal experiments; C.H. and S.G wrote python script for *in vitro* and *ex vivo* experiments; K.R. ensured collaboration between MUMC+ and Maastricht University for tissue collection; A.W and S.H performed electrophysiology experiments; G.K. supervised the stem cell experiments; H.C., Y.T., D.G., and S.H. supervised all aspects of the work.

## Data availability

Source data are provided with this paper.

## Code availability

The code for neuronal stimulation and fluorescence image analysis is provided with this paper.

---

**Figure S1.**
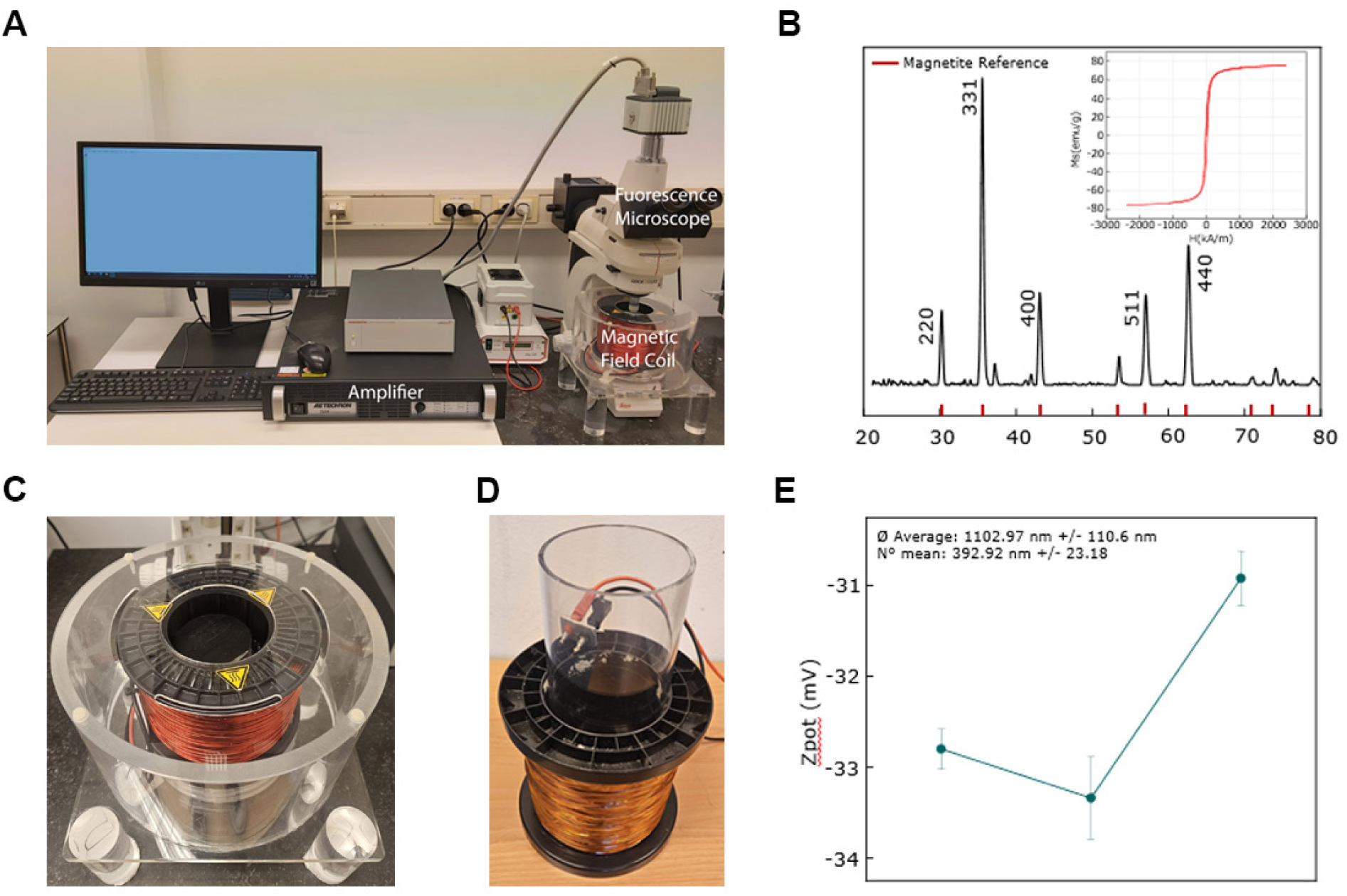
Magnetomechanical setup and MNDs characterisation. **A.** Overview of in vitro and ex vivo magnetomechanical stimulation setup. **B.** XRD spectrum and VSM analysis of the MNDs confirm the crystalline phase and magnetic saturation of magnetite. **C.** Overview of in vivo magnetomechanical stimulation setup. **D.** Zeta potential and average diameter of MNDs after PMAO coating.

**Figure S2.**
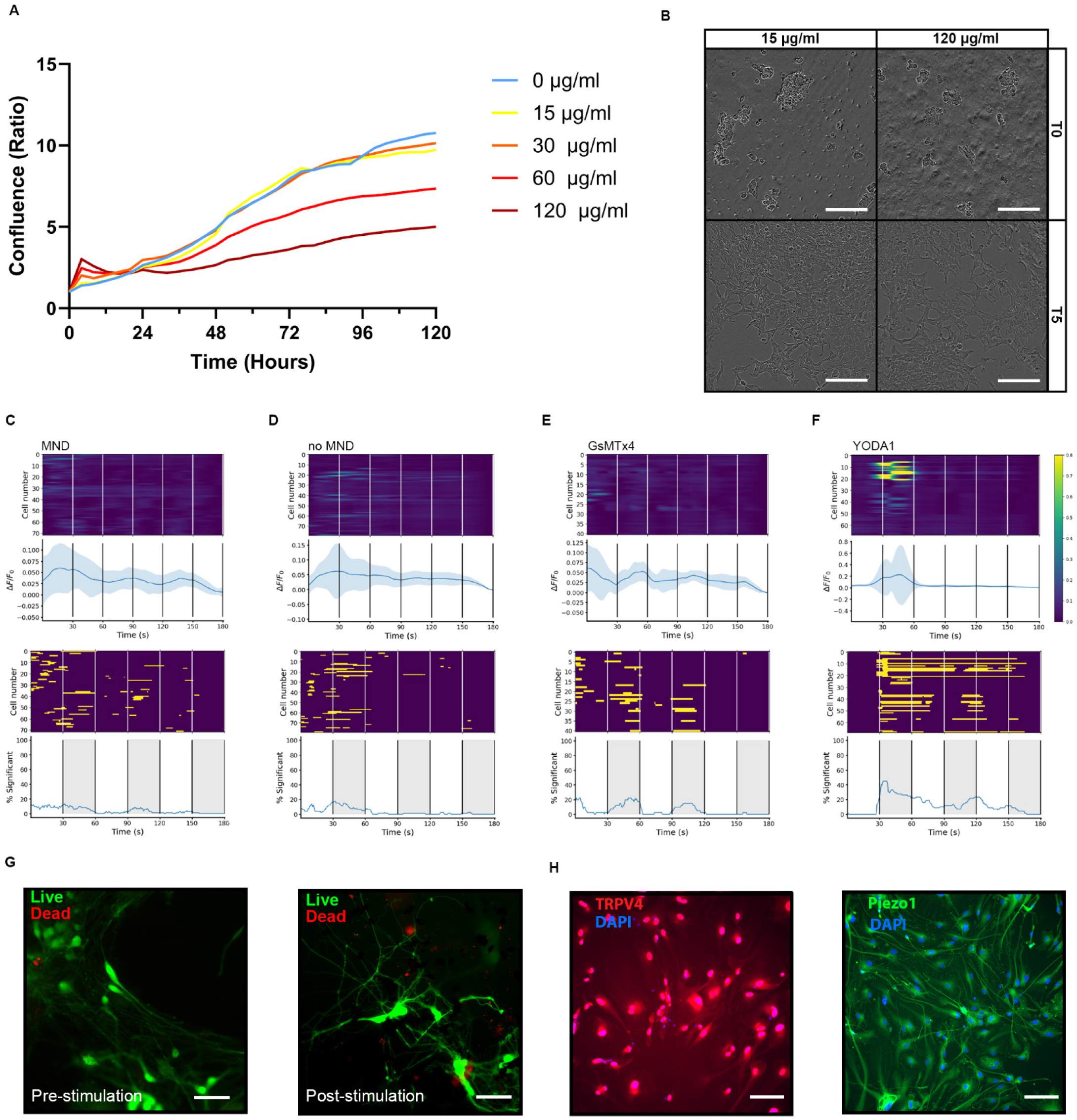
In vitro magnetomechanical control. **A.** Real-time quantification of cell death in HEK293 cells following 5 days of exposure to MNDs at various concentrations was performed in triplicate. The concentration in all further in vitro experiments was 30 µg/ml. **B.** Images of cells immediately after exposure and 5 days of exposure to MNDs at 15 µg/ml and 120 µg/ml. Scale bar = 150 µm. **C-D** Heatmaps (top) and significance plots (bottom) of fluorescence intensity changes recorded from Fluo-4 AM transients observed in Piezo1 and TRPV4 expressing hESC-derived neurons during magnetic field stimulus with (C) and without (D) MNDs during AMF Off. **E.** Magnetomechanical cation channel inhibition of Piezo1 and TRPV4 through GsMTx4. **F.** Pharmacological activation of Piezo1 through YODA1. Each condition was repeated across six different cultures, with ∼10 cells randomly selected for analysis. **G.** Confocal images of hESC-derived neurons loaded with the calcium indicator Fluo-4 AM and propidium iodide (PI) for live/dead staining before and after mechanostimulation. Fluo-4 AM stains live cells in green, and PI stains dead cells in red. Scale bar = 50 µm. **H.** Confocal microscope images of hESC-derived neurons stained for TRPV4 (red), Piezo1 (green) and DAPI (blue). Scale bar = 50 µm.

**Figure S3.**
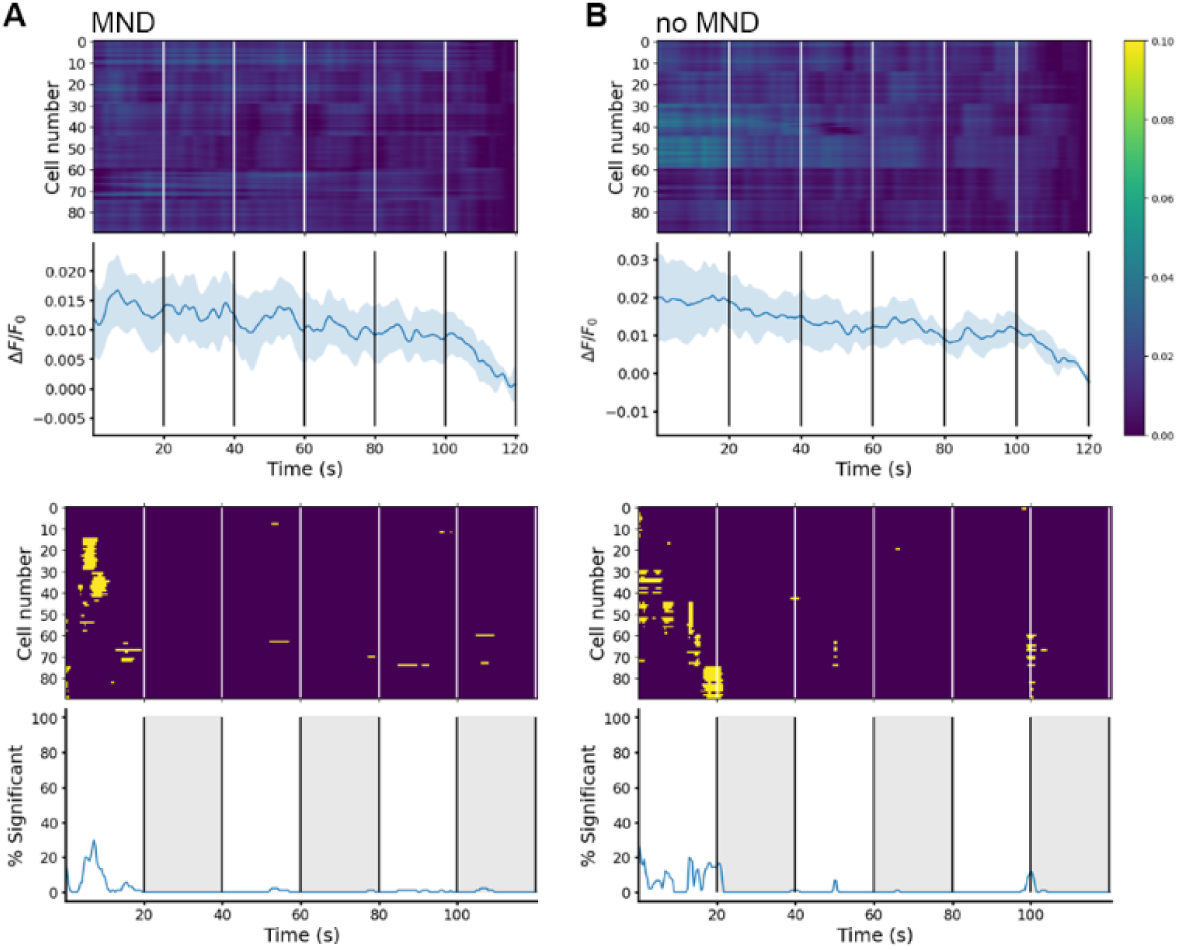
Ex vivo magnetomechanical control. Heatmaps (top) and significance plots (bottom) of fluorescence intensity changes recorded from Fluo-4 AM transients observed in Piezo1 and TRPV4 expressing hBSC during magnetic field stimulus with (A) and without (B) MNDs during AMF Off. Each condition was repeated across six different cultures, with ∼15 cells randomly selected for analysis.

**Figure S4.**
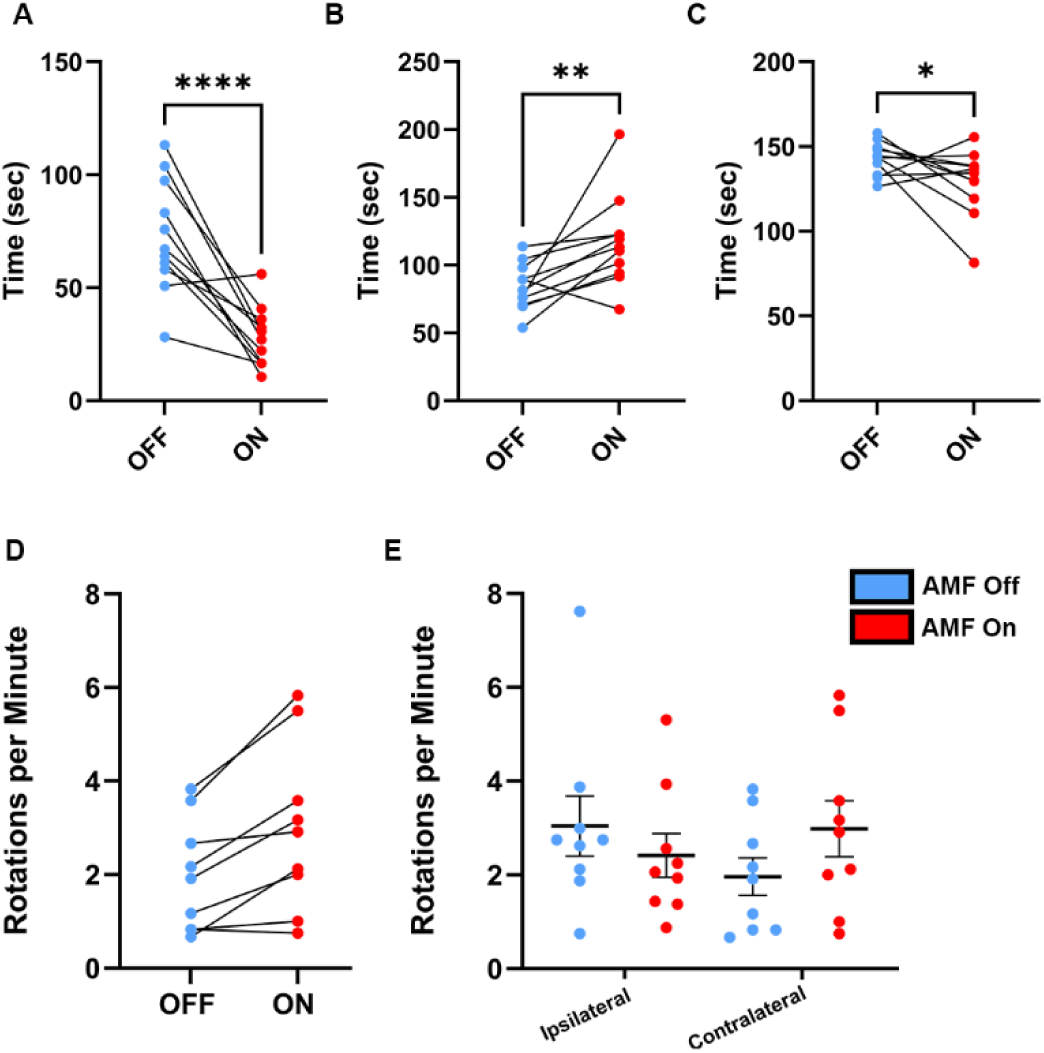
Remote magnetomechanical neuromodulation of mouse behaviour. **A.** Decreased total time spent in the centre (p=0.0002). **B.** Increased total time spent in the corners (p=0.0059). **C.** Decreased total time spent at the walls (p=0.0407) during the 5-minute OFT. **D.** Contralateral rotations of STN mDBS (n=11) during AMF On and Off conditions. **E.** Rotational behaviour presented in ipsilateral and contralateral rotations of STN mDBS (n=11) during AMF On and Off. * Indicates p < 0.05, ** < 0.01, **** < 0.0001. One-tailed t-test, data presented as mean ± S.E.M.

**Figure S5.**
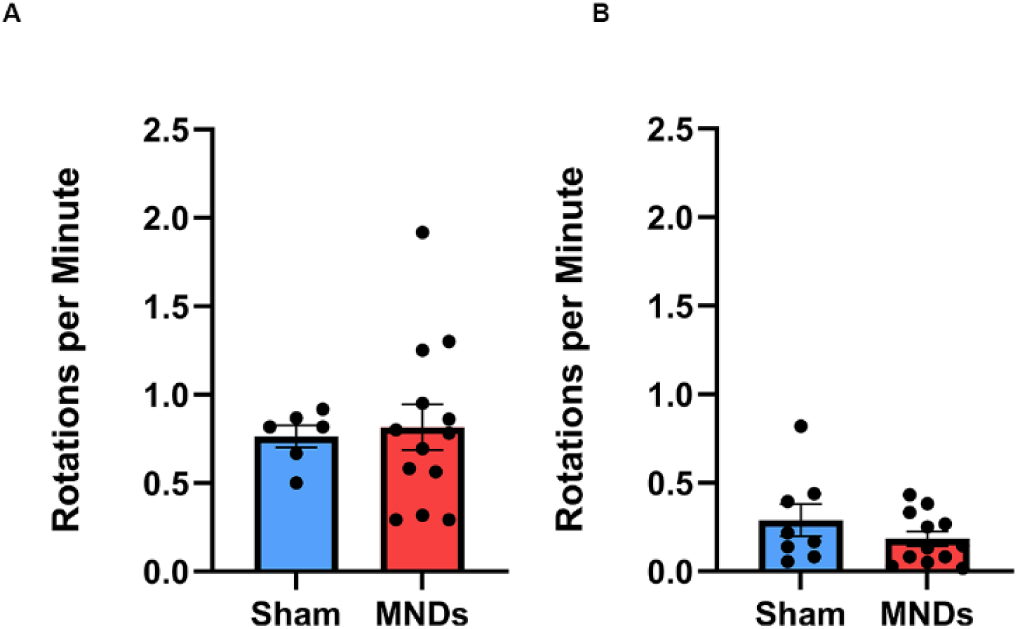
The effects of magnetomechanical STN DBS on rotational behaviour. **A.** Ipsilateral rotations per minute. **B.** Contralateral rotations per minute of the sham (n=8) and STN mDBS (n=13). Data presented as mean ± S.E.M.

**Figure S6.**
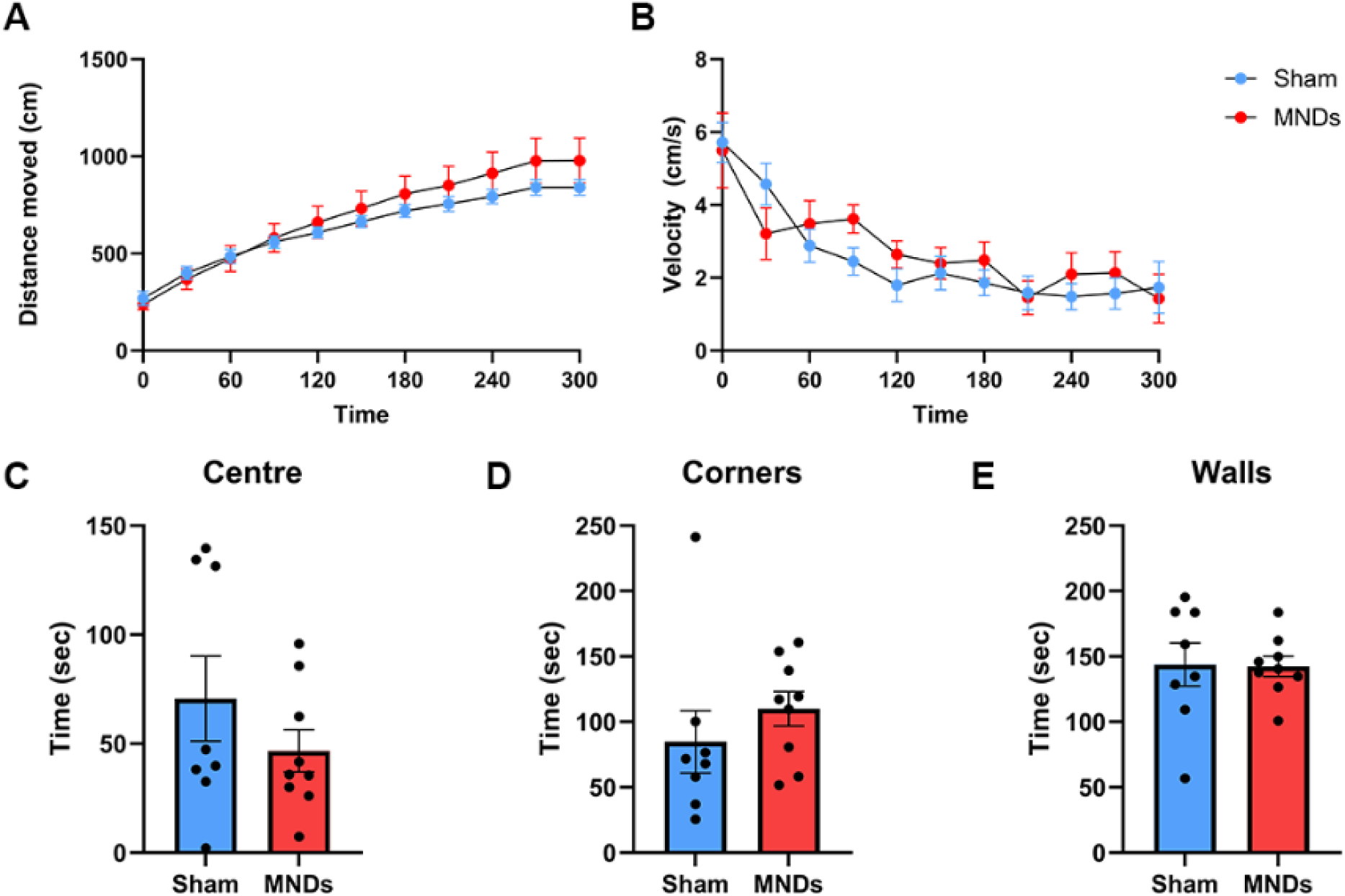
The effects of magnetomechanical STN DBS during the Open Field Test (OFT). **A.** Total distance moved (in cm) in the OFT ± S.E.M. for sham (n=7) and STN mDBS (n=9) mice. There was a non-significant increase between the groups when considering 30-second time bins. **B.** Velocity (in cm/sec) in the OFT ± S.E.M. for sham (n=7) and STN mDBS (n=9) mice. There was a non-significant increase between the groups when considering 30-second time bins. **C.** Total time spent in the centre. **D.** Total time spent in the corners. **E.** Total time spent at the walls during the 5-minute OFT. Data presented as mean ± S.E.M.

**Figure S7.**
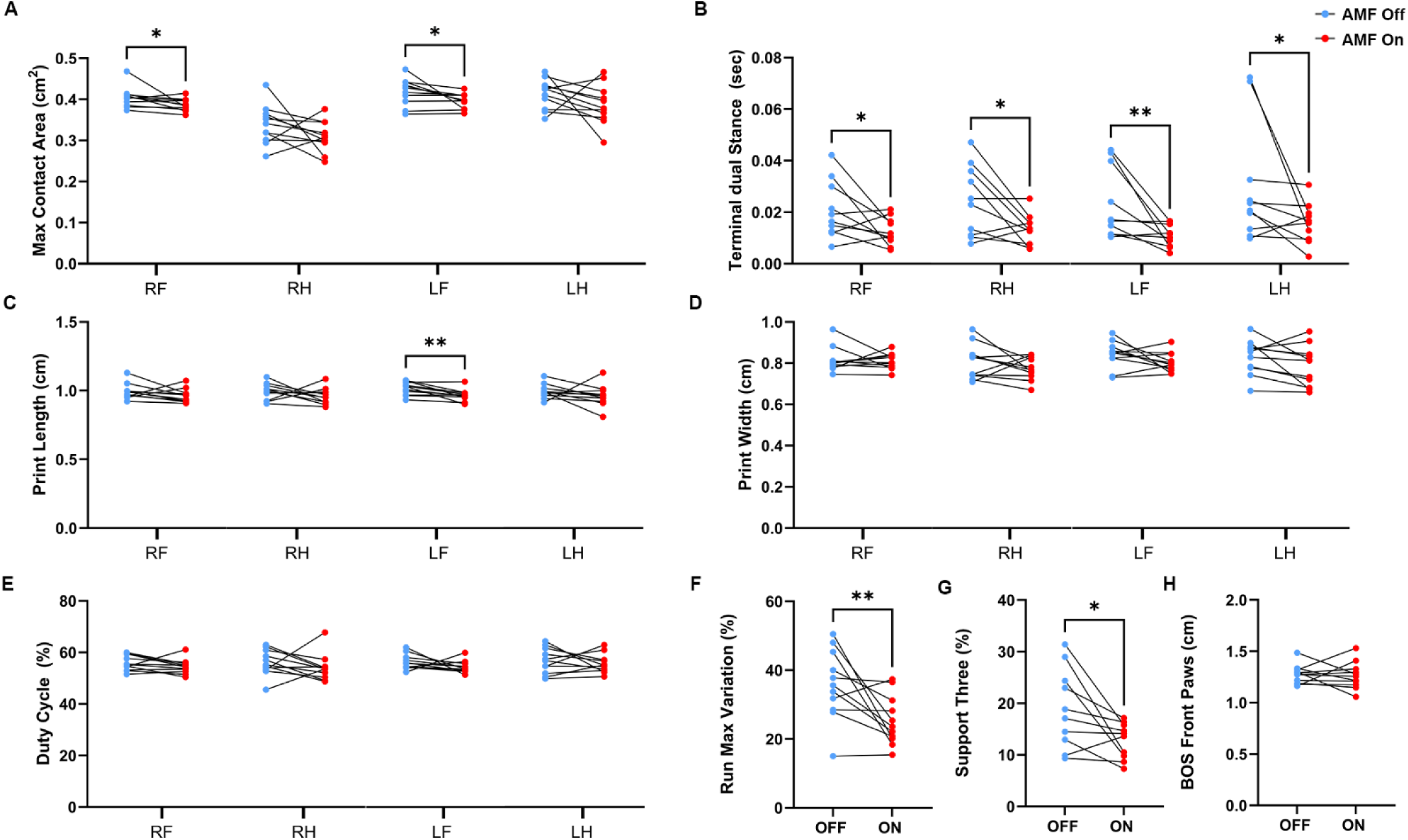
Magnetomechanical STN DBS improves motor behaviour in naïve male mice. Representative CatWalk XT results of four runs. AMF stimulation resulted in increased max contact area (p ≤ 0.0255) **(A)**, a decrease in terminal dual stance (p ≤ 0.0425) **(B)**, decreased print length (p = 0.0026) **(C)**, no change in print width (P ≤ 0.0358) **(D)**, no change in speed duty cycle **(E)**, a decrease in run max variation (p = 0.0075) **(F)**, a decrease in support three (p = 0.0134) **(G)**, and a no change in base of support front paws **(H)** between AMF On and AMF Off (n=11) as revealed by an independent one-tailed t-test. * Indicates p < 0.05, ** < 0.01. Data presented as mean ± S.E.M. RF: right front paw, RH: right hind paw, LF: left front paw, LH: left hind paw.

**Figure S8.**
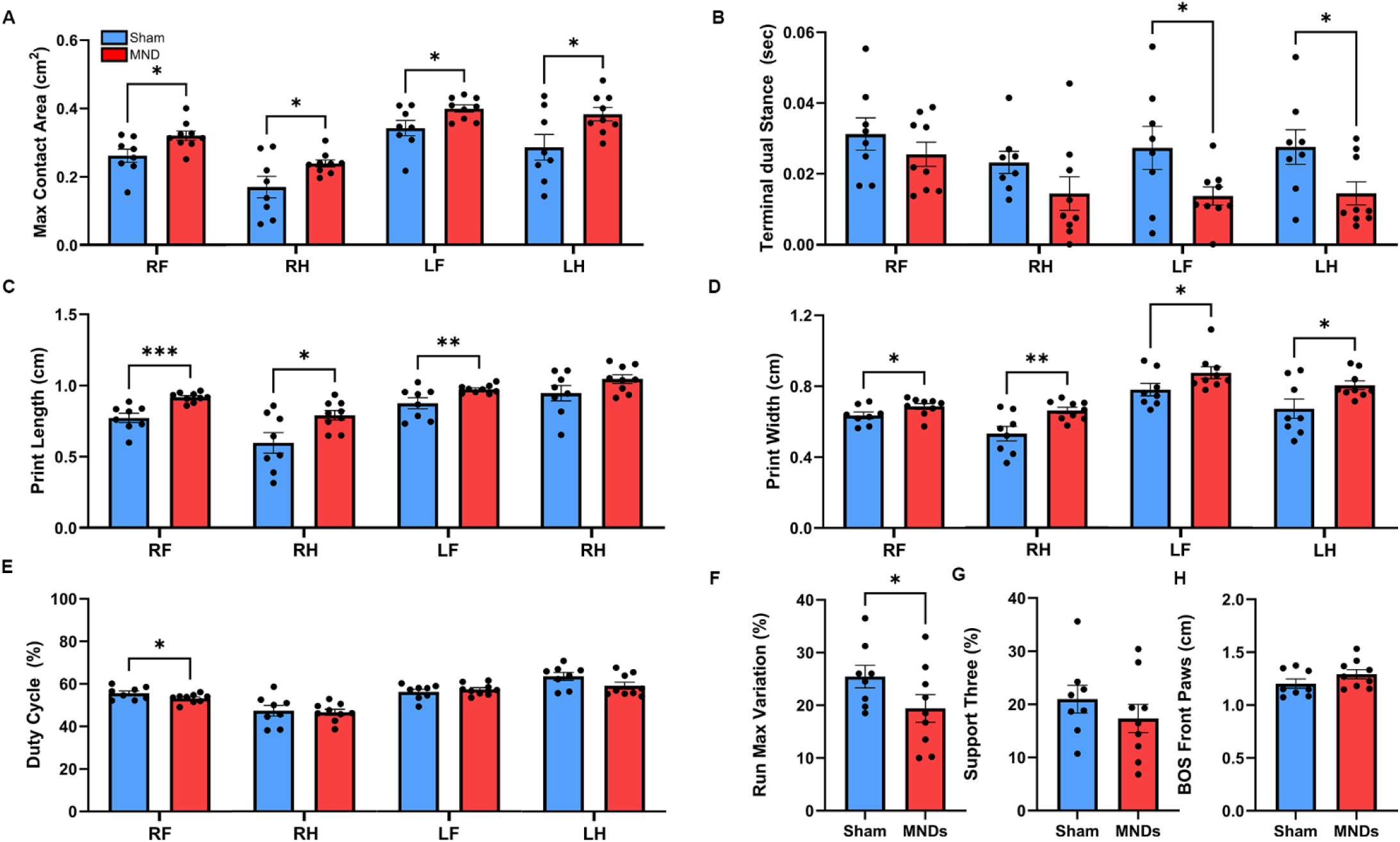
Magnetomechanical STN DBS alleviates severe Parkinsonian symptoms. Representative CatWalk XT results of four runs. AMF stimulation resulted in increased max contact area (p≤0.0228) **(A)**, a left-sided decrease in terminal dual stance (p ≤ 0.0246) **(B)**, increased print length (p ≤ 0.0110) **(C)**, increased print width (P ≤ 0.0358) **(D)**, decreased speed duty cycle (p = 0.0352) **(E)**, decreased run max variation (p = 0.0485) **(F)**, no change in support three **(G)**, no change in base of support of front paws **(H)** between sham (n=9) and STN mDBS (n=8) as revealed by an independent one-tailed t-test. * Indicates p < 0.05, ** p < 0.01, *** p < 0.001. Data presented as mean ± S.E.M. RF: right front paw, RH: right hind paw, LF: left front paw, LH: left hind paw.

